# Mixed selectivity coding of sensory and motor social signals in the thalamus of a weakly electric fish

**DOI:** 10.1101/2021.04.04.438391

**Authors:** Avner Wallach, Alexandre Melanson, André Longtin, Leonard Maler

## Abstract

Recent studies have shown that high-level neural activity often exhibits mixed selectivity to multivariate signals. How such representations arise and how they modulate natural behavior is poorly understood. The social behavior of weakly electric fish is relatively low-dimensional and easily reproduced in the laboratory. Here we show how electrosensory signals related to courtship and rivalry in *Apteronotus leptorhynchus* are represented in the preglomerular nucleus, the thalamic region exclusively connecting the midbrain with the pallium. We show that preglomerular cells convert their midbrain inputs into a mixed selectivity code that includes corollary discharge of outgoing communication signals. We discuss how the preglomerular pallial targets might use these inputs to control social behavior and determine dominance in male-male competition and female mate selection during courtship. Our results showcase the potential of the electrocommunication system as an accessible model for studying the neural substrates of social behavior and principles of multi-dimensional neural representation.

## Introduction

In mammals, sensory input can generate rich representations of complex events that occur at particular locations and times. These representations can be stored and recalled to guide ongoing behavior. How are complex multi-dimensional sensory inputs and motor outputs combined to enable representations that bind different sensory-motor attributes into meaningful wholes? Recent studies suggest that combined encoding of multiple stimulus features, i.e., neurons with mixed selectivity, serve such a role (Rigotti et al., 2013). Nonlinear mixed selectivity neurons have further been hypothesized to form the bases of a high dimensional coding space (Rigotti et al., 2013) that supports robust computations (Johnston et al., 2020). Social behavior of weakly electric fish offers an attractive model for connecting mixed selectivity coding of sensory input to ethologically meaningful representations that guide behavior. The electrocommunication signals associated with social behavior are relatively low dimensional, i.e. they can be described using few variables, and are easy to record and mimic in the laboratory. In this paper, we describe the responses of neurons in a thalamus-like structure (Wallach et al., 2018), the preglomerular nucleus (PG), to the well-defined electrocommunication signals of the electric fish, *Apteronotus leptorhynchus*.

Electrocommunication signals of Apteronotus have been described in laboratory settings (K. D. Dunlap & J. Larkins-Ford, 2003; Engler & Zupanc, 2001; Hagedorn & Heiligenberg, 1985; Hupe & Lewis, 2008; Walz et al., 2012; Zakon et al., 2002; Zupanc, 2002) and in the fish’s natural environment (Henninger et al., 2018). The motor pathways that generate electrocommunication signals (Metzner, 1999), and the responses of electroreceptors, medullary and midbrain electrosensory neurons to such signals (Chacron et al., 2011; Clarke et al., 2015; Krahe & Maler, 2014; Marsat et al., 2012) have been thoroughly investigated. In this paper, we show that PG neurons encode various combinations of motor and sensory attributes of electrocommunication signals and link such combinatorial coding to the natural social interactions of Apteronotid fish. We further discuss how such coding might be used by the dorsal telencephalon (pallium) to generate ethologically meaningful representations of the social interactions. In particular, we emphasize that the responses of PG neurons exhibit ‘mixed selectivity’ to the electrocommunication signals emitted during natural behavior and can therefore be considered as bases of a high-dimensional neural coding space instantiated in pallium.

### Components of electric social behavior

South American gymnotiform fish include genera (Apteronotids and Eigenmannia) that emit high frequency sinusoidal electric organ discharges (EOD) and have electroreceptors tuned to their EOD; this combination permits these nocturnal fish to navigate, hunt and communicate in the dark (Chacron et al., 2011; Clarke et al., 2015; Jun et al., 2016; Marsat et al., 2012). Each individual fish has a characteristic EOD frequency. The distribution of these frequencies is sexually dimorphic: in A. leptorhynchus males use frequencies in a high range (750-1100Hz) whereas females use a low range (600-750Hz) (Zakon et al., 2002). Furthermore, male dominance is signaled by a fish’s EOD frequency, with higher frequencies indicative of more dominant males (Fugere et al., 2011; Zakon et al., 2002). When two such fish are in proximity, their waveforms interfere so as to create amplitude and phase modulations of the EOD referred to as a “beat”. The beat frequency is equal to the difference of their EOD frequencies (denoted ΔF), e.g., if two males with frequencies of 918 Hz and 900 Hz are in proximity to one another, an 18 Hz beat will be generated (**Fig. 1**). The beat frequency, therefore, informs each fish about the sexual identity and dominance of the other; different-sex interactions usually result in high frequency beats (**Fig. 1A**), while same-sex interactions result in lower frequency beats (**Fig. 1B**). Additionally, the precise beat frequency may be used to distinguish between specific individuals, and fish can indeed learn to precisely identify beat frequencies (Harvey-Girard et al., 2010).

**Figure 1:**
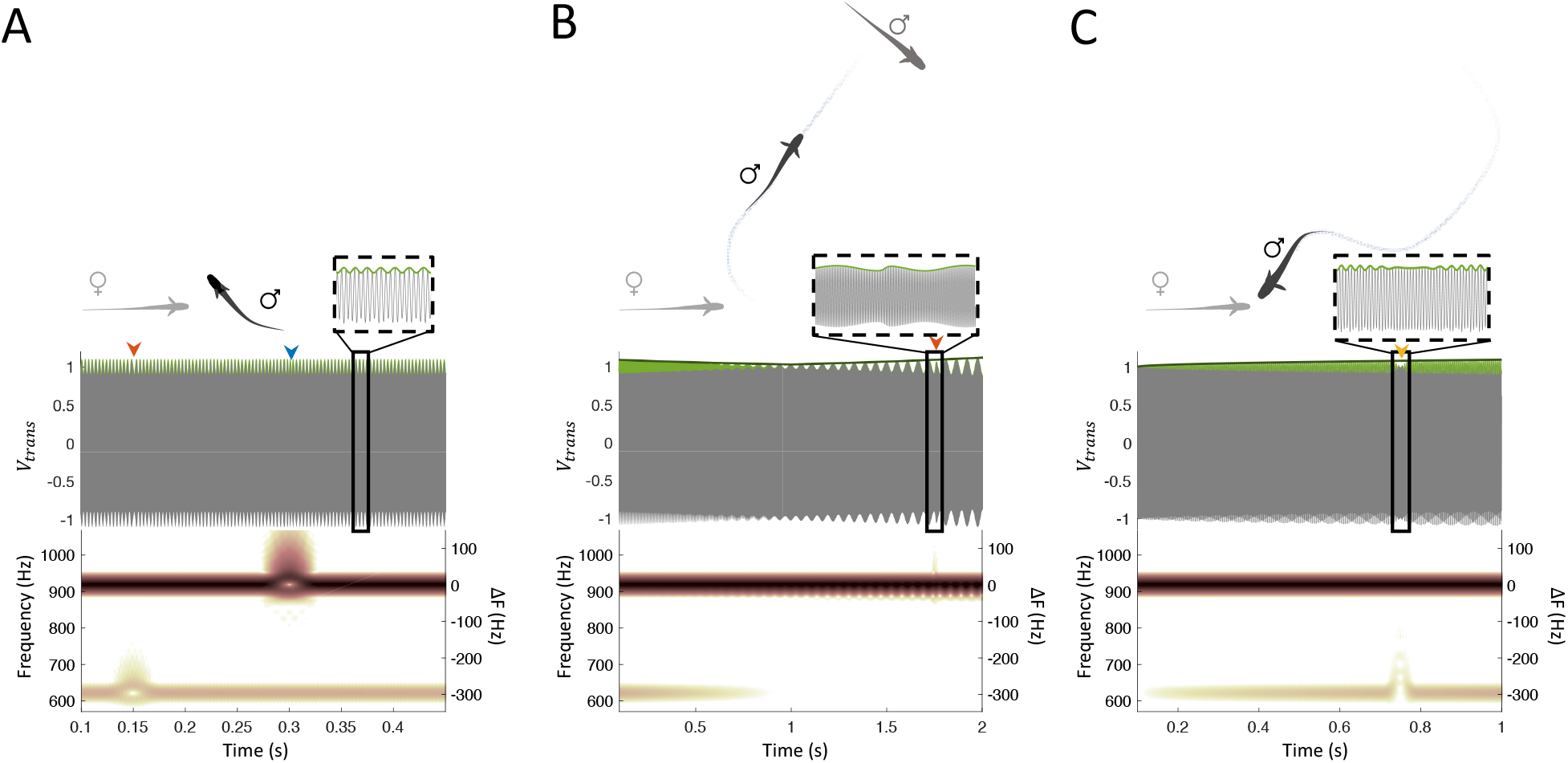
Components of electric social behavior. Each individual fish has a characteristic EOD frequency; male frequency is typically in the high range (>750Hz) while that of females is in the low range (<750Hz). Top: illustration of different social scenarios; Middle: transdermal potential sensed by the male fish (black); Bottom: Spectrogram of the potential. (**A**) A male fish (918Hz) courting a female (621Hz). The interference between the two EODs creates a modulation pattern called a *beat* (green), whose frequency equals the absolute difference between the two EOD frequencies (i.e., 297Hz). The female emits a *small chirp* (red arrowhead), which the male reciprocates with small latency (blue arrowhead). (**B**) The courting male detects an approaching rival male (900Hz) and turns to drive it away. The relative motion alters the *envelope* of the beat patterns (dark green line) – *the* female induced high-frequency beat subsides while the low-frequency male induced one increases. The rival male emits a *small chirp* to signal submission (red arrowhead). (**C**) The male returns to the female, who emits a *long chirp* to signal spawning (yellow arrowhead).

Electroreceptors and a subset of brainstem electrosensory neurons are tuned to beats over the frequency range that occurs during natural behaviors (**Fig.1**; **Fig. 2A**) (Chacron et al., 2005; Krahe & Maler, 2014; Savard et al., 2011; Vonderschen & Chacron, 2011). The amplitude of the beat (depth of modulation) is strongly affected by the distance between the two fish and by their relative orientation. When the two fish move towards (away from) one another, the amplitude increases (decreases), thus generating a ‘second order’ envelope signal (**Fig. 1B)** (Fotowat et al., 2013; Metzen & Chacron, 2019; Yu et al., 2012; Yu et al., 2019). Therefore, this envelope signal conveys information about the relative motion of a conspecific, e.g. its approach (retreat). A subset of electroreceptors and their target medullary and midbrain electrosensory neurons respond to envelope signals (**Fig. 2A**) (Metzen & Chacron, 2019).

**Figure 2:**
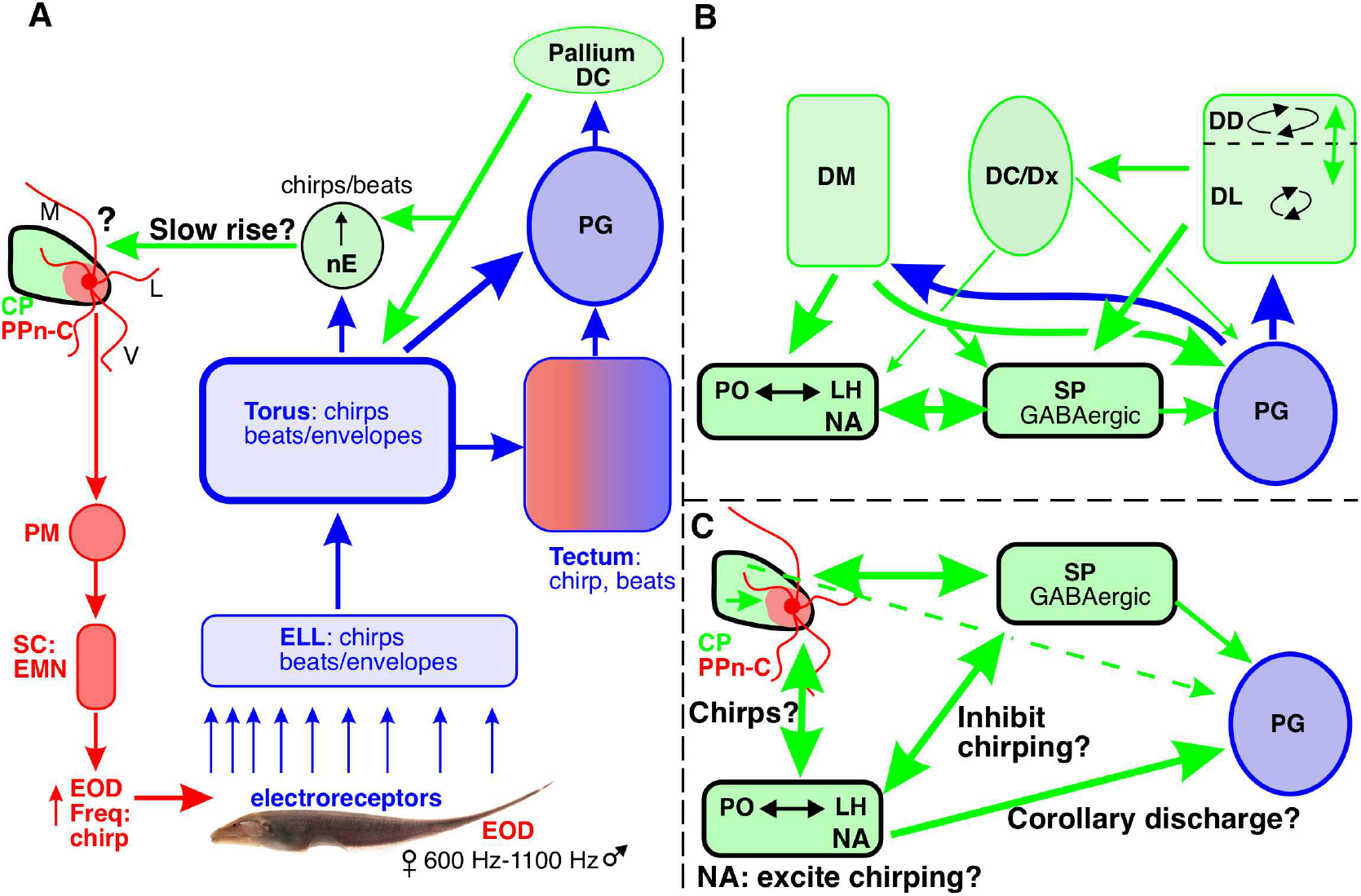
The anatomical substrates of electrocommunication during social behavior. Red nuclei and arrows indicate the electromotor pathways. Blue nuclei and arrows indicate the ascending electrosensory pathways. Green nuclei and arrows illustrate intermediate bran structures and pathways that, because of their connectivity and/or physiology, appear to link high-level processing of electrocommunication signals and control of chirping. Feedback pathways to the electrosensory lobe (ELL) and minor connections are not illustrated. Text followed by a question mark highlights hypotheses on function made in the Discussion, e.g., “Corollary discharge?”. **(A**) *Electromotor pathways*. A medullary pacemaker (PM) drives spinal cord (SC) electromotor neurons (EMN). The axons of electromotor neurons constitute the neurogenic electric organ that generates the electric organ discharge (EOD). A diencephalic prepacemaker nucleus (PPn) located lateral to the central posterior nucleus (CP) projects to the PM and can modulate pacemaker frequency. A ventro-lateral magnocellular component of the PPn, PPn-C, can transiently increase the EOD frequency and thus generate electrocommunication signals – chirps. Discharge of CP cells medial to the ‘chirp’ cells will evoke gradual increases in EOD frequency (hence PPn-G, not illustrated). PPn-C cells have sparse medial (M) and lateral (L) and very extensively ramifying ventral (V) dendrites. Medial dendrites penetrate CP and also ramify in the neuropil dorsal to CP. *Ascending electrocommunication pathways*. The electric field generated by the EOD stimulates the array of cutaneous electroreceptors. These project to the ELL which, in turn, projects to the midbrain torus semicircularis (TS). Most neurons in ELL and a subset of TS neurons respond to beats, envelopes and chirps. TS neurons project to tectum, n. electrosensorius (nE) and the preglomerular nucleus (PG); descending TS projections are not illustrated. A subset of tectal neurons respond to beats and chirps (Heiligenberg & Rose, 1987); envelope signals were not used in studies of tectum. The tectum is colored red and blue to indicate that it has both ascending and descending projections. nE is a complicated structure with numerous subdivisions; cells within some subdivisions are responsive to beats and chirps; envelope stimuli were not used in studies of nE. A portion of nE (nE up or nE↑) projects to CP and to the lateral and medial dendritic territories of PPn-C neurons. This projection likely mediates the slow rise electrocommunication signal (Heiligenberg et al., 1996). PG projects to dorsolateral pallium (DL, see Panel B). **(A – cont**.) DL in turn projects to the central pallium (DC, Panel C). DC can, via a strong projection to TS, modulate the ascending electrosensory input to PG. In addition, DC provides a strong and specific projection to the nE↑ subdivision of nE (Giassi, Duarte, et al., 2012). This projection possibly mediates pallial control of slow rises. **(B**) PG projects primarily to the dorsolateral (DL) and dorsomedial (DM) pallium; curved black arrows indicate local recurrent circuitry in DL (Trinh et al., 2016) and global recurrent circuitry in DD (Elliott et al., 2017). DL and the interconnected dorsal pallium (DD) can potentially influence processing and/or control of electrocommunication via strong projections to the SP and minor projections, via central pallium (DC,Dx), to the lateral hypothalamus (LH) and PG. DM has strong descending projections to LH and subpallium as well as feedback projections to PG. **(C)** The subpallium (SP), preoptic region (PO), LH, CP/PPn-C are all reciprocally interconnected. SP projecting to PG, PO, CP/PPn-C and LH is now known to be GABAergic and its projection to PPn-C and LH may inhibit chirping, and thus cause the post long chirp reduction in small chirp emission (Henninger et al., 2018). LH peptidergic neurons project strongly to the ventral dendrites of PPn-C neurons; glutamate application to these dendrites evokes small chirps (Kawasaki et al., 1988). LH is strongly innervated by noradrenergic afferents (NA) (Sas et al., 1990) and might be the site at which NA excites chirping (Maler & Ellis, 1987). The preoptic projections to CP/PPn include cells expressing vasotocin (Pouso et al., 2017) and choline acetyltransferase (Toscano-Marquez et al., 2020); this projection may drive long chirps (Bastian et al., 2001). It is notable that LH and the preoptic area provide strong input to PG that we hypothesize to mediate chirp corollary discharge. CP provides weak input (dashed arrow) to PG and may also project to the PPn-C medial dendrites (Wong, 1997a). It is also important to note that PPn-C does not project to PG (Giassi, Duarte, et al., 2012).

Apteronotid fish communicate by increasing their EOD frequency during same and opposite sex social interactions (**Fig. 1**) (Hupe & Lewis, 2008; Walz et al., 2012). They do so by either transiently increasing their EOD frequency (chirps) or by more gradual and sustained EOD frequency increases (slow rises) (Zakon et al., 2002; Zupanc, 2002). In this paper we focus on chirps because the responses to such signals have been well characterized in the ascending electrosensory pathway and no equivalent studies have been carried out for slow rises (but see **Fig. 2A**). Two major classes of chirps have been reported in laboratory studies (Zakon et al., 2002; Zupanc, 2002) and in the wild (Henninger et al., 2018) and they have been simply described as ‘small’ and ‘long’ chirps. Small chirps are most common and can occur during both male-male and male-female interactions (**Fig. 1A and B)** (Zakon et al., 2002; Zupanc, 2002). Long chirps typically occur during opposite sex interactions (Bastian et al., 2001; Hagedorn & Heiligenberg, 1985; Henninger et al., 2018) and, especially, during courtship (**Fig. 1C)** (Hagedorn & Heiligenberg, 1985; Henninger et al., 2018). Electroreceptor, medullary and midbrain neuron responses to both small and long chirps have been described (Benda et al., 2005, 2006; Marsat & Maler, 2010; Marsat et al., 2009; Vonderschen & Chacron, 2011; Walz et al., 2012). Medullary neurons respond in a similar manner to the fish’s own small chirp and to a mimic of another fish’s small chirp (Marsat et al., 2009), and it is therefore not clear how a fish can distinguish self from conspecific chirps. In summary, electric social behavior is composed of several features: beats, whose frequency conveys fish identity, envelopes that convey relative motion and at least three types of social signals: discrete stereotypic transient increases in EOD frequency (small and long chirps) and gradual sustained small increases in EOD frequency (slow rises). Here we focus on PG neuron responses to beats, envelopes and chirps.

### The anatomical substrate of electric social behavior

In **Fig. 2A** we sketch out the electrosensory circuitry by which beats, envelopes and chirps are transmitted from electroreceptors to the midbrain and diencephalon (Bell & Maler, 2005; Carr et al., 1981; Carr et al., 1982; Giassi, Duarte, et al., 2012), and the motor pathways known to control the emission of chirps (Metzner, 1999). Early studies suggested interconnections between chirp detection and emission were confined to the midbrain and diencephalon (Keller et al., 1990; Walz et al., 2012), and failed to find direct ascending input to the preglomerular nucleus and pallium (Carr et al., 1981). In **Fig. 2B and C** we illustrate circuitry from more recent studies that connect PG representation of electrocommunication signals to higher level pallial processing and reveal intersecting and nested loops connecting chirp/beat/envelope detection and chirp emission. **Panels A and B** also illustrate routes by which pallium can modulate its own input via descending pathways to the midbrain and PG. Below, we describe the ‘mixed selectivity’ of PG neurons to various combinations of beats, envelopes, and self and exogenous chirps, and possible responses to chirp emission corollary discharge. We suggest that the pallial targets of PG can combine these inputs to generate rich representations of social behavior as neural activity trajectories in a high dimensional coding space.

## Results

We recorded from N=98 electrocommunication responsive PG neurons in N=26 fish. Recordings from PG were done as previously described (Wallach et al., 2018). Most cells were found in recording sites that were about 200µm medial to those in that study and were therefore within the medial PGm (Giassi, Duarte, et al., 2012). As was the case in our study of motion coding responses in PG (Wallach et al., 2018), we attempted to sample broadly within PGm along both dorso-ventral and rostro-caudal dimensions.

Our stimulus set included sine waves that mimicked the EOD of a conspecific Apteronotus. These stimuli produce beats at the difference frequency (ΔF) between the EOD mimic and the fish’s own EOD frequency. A wide range of ΔFs was used to mimic the signals produced by fish with widely varying EOD frequencies. We also modulated the beat amplitude over longer time scales to produce low frequency envelopes that occur during movement of the conspecific towards or away from the experimental fish. Finally, we transiently modulated the frequency and duration of the stimulus sine wave to produce mimics of small and long chirps; we refer to these as exogenous small and long chirps. **Figure 1** illustrates the stimuli used and full experimental details are presented in the Methods section. In this paper we start with describing PG neuron responses to self and exogenous chirps. We then describe the responses to beats and envelopes. Finally, we describe the responses of PG neurons to combinations of various types of chirps, beats and envelopes.

Some fish (N=11) emitted small chirps (‘self-chirps’) when presented with an EOD mimic and PG cell spiking responses to small self-chirps were also recorded (N=21 cells). No fish ever emitted a long chirp in our recording set-up, and we therefore have no data on the responses of PG cells to long self-chirps. Self-chirps were comparable to small chirps as previously reported (Marsat et al., 2009) with frequency excursions of ∼100 Hz and durations of 10-20 ms (**Fig. 3A**). The exogenous small chirps matched these values (**Fig. 3B**) and the exogenous long chirp stimuli were designed to conform to their descriptions in the literature (**Fig. 3C**) (Bastian et al., 2001; Engler et al., 2000; Engler & Zupanc, 2001; Henninger et al., 2018). PG neurons typically exhibited a low level (4.1±4.6 Hz) of bursty baseline activity similar to that previously reported for lateral PG cells (Wallach et al., 2018). The beat stimuli increased the discharge of many, but not all, chirp responsive PG neurons; details are provided where appropriate. The latency of the first spike response to a self or exogenous chirp was a critical measurement and was therefore rigorously and conservatively estimated to be certain that such spikes were in response to a chirp and not baseline discharge or spikes evoked by the beat stimulus (see Methods). Response durations are calculated from the time interval between estimated first to last spike.

**Figure 3:**
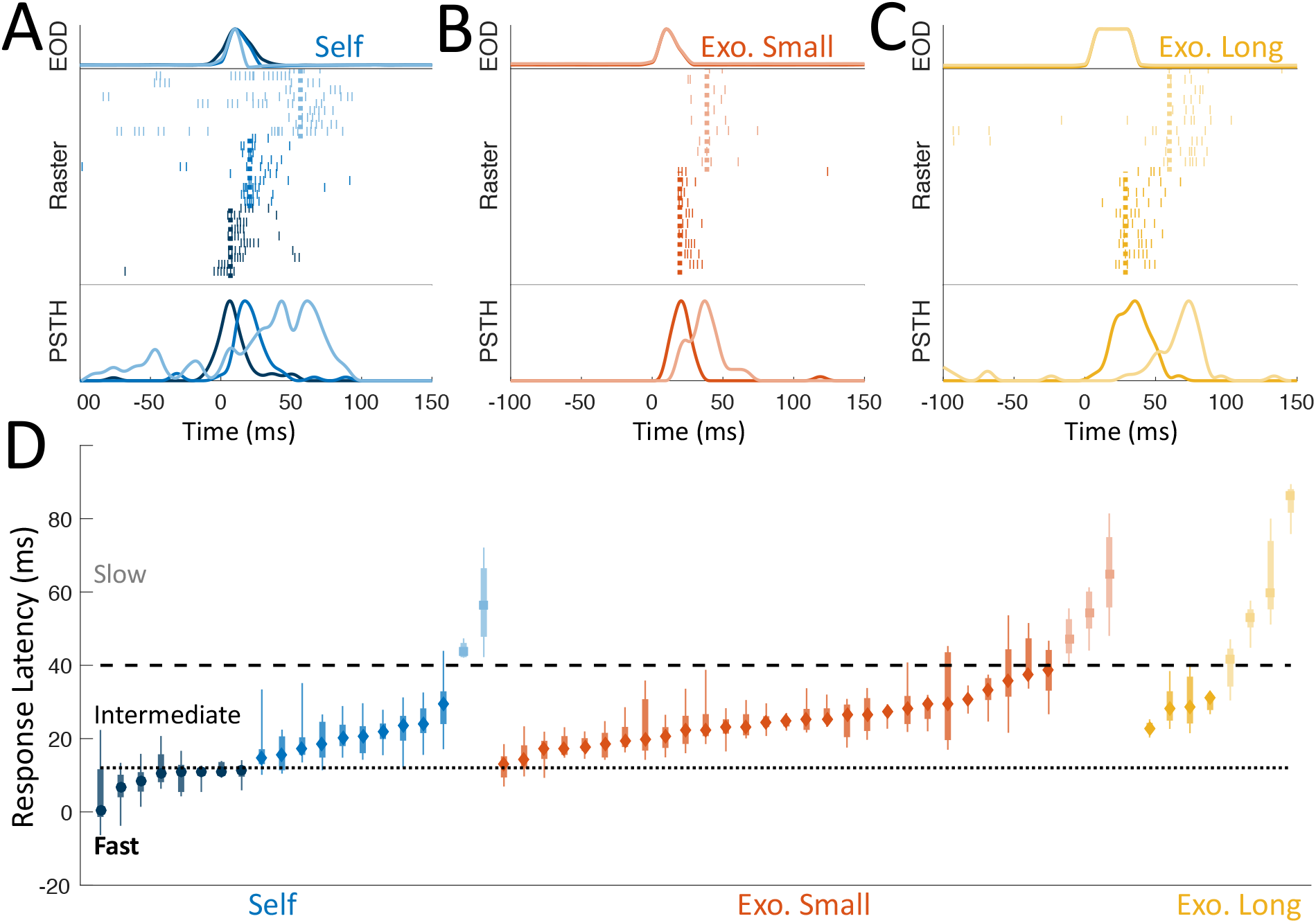
PG cells respond to different chirp types with various latencies. (**A-C**) Top: EOD rate excursion (normalized); Middle: raster plot depicting spike responses to individual chirp instances (dashed lines-median first spike latency); Bottom: PSTH across chirps. (**A**) Three different PG cells responding to self-generated small chirps with small (dark blue), intermediate (blue) and slow (light blue) latencies. (**B**) Two cells responding to exogenous small chirp with intermediate (red) and slow (light red) latencies. (**C**) Two cells responding to exogenous long chirp with intermediate (yellow) and slow (light yellow) latencies. (**D**) Latencies of all PG responses to the different chirp types. Circles/diamonds/squares: fast/intermediate/slow median latencies, respectively; thick lines: Inter-quartile range (IQR); thin lines: 10%-90% range.

The anatomy of ascending electrosensory pathways to PG (illustrated in **Fig. 2A and B**) leads to our naïve initial assumption that the responses of PG neurons to electrocommunication signals would be similar to those already reported in hindbrain (ELL) and midbrain (TS), but with a slightly longer latency. We present these predictions because they highlight striking discrepancies from our data and lead to a novel view of electrocommunication signal coding and its role in social behavior.

### Responses to self vs exogenous chirps

ELL neurons respond at the same short latency to small self- and exogenous chirps with spiking confined to a short burst that does not outlast the chirp (Marsat et al., 2009). TS neurons respond to small exogenous chirps in a similar manner, emitting a short burst within a narrow fast latency range (Vonderschen & Chacron, 2011).

### Responses to small versus long chirps

Different ELL neurons respond to small versus long chirps but the responses have similar latencies (Marsat & Maler, 2010; Marsat et al., 2009). TS neurons respond in a similar manner to both types of chirps – with a short burst at the same latency; there are neurons that maintain response selectivity while others respond to both chirp types (Vonderschen & Chacron, 2011).

#### Chirp response latency distributions suggest possible PG cell categories

PG neuron responses to self (**Fig. 3A**), exogenous small (**Fig. 3B**) and long (**Fig. 3C**) chirps extend over a very wide range of latencies and spiking statistics (see **Table 3S1** for all latency values). Comparisons of responses can be fruitfully made both within and across chirp types. The most striking parcellation of chirp responses is based on response latencies from chirp onset (**Fig. 3D**) and we use the resulting possible categories as a framework for more detailed analyses of chirp spiking responses. Median response latencies for a subset of self-chirp responsive cells are <12 ms while all responses to exogenous small chirps have latencies ≥ 12 ms, and the fastest responses to long chirps have latencies >22 ms. We therefore define a “fast latency” category unique to self-chirps with median response latencies ranging from 0.4 to 11.3 ms (N=8). We note that the first evoked spikes within this category are smaller and range from −6.3 to 9.32 ms. Given that self-chirp durations were 10-20 ms, the first spikes of fast responses were typically initiated before or soon after chirp onset. (**Figs. 3A, 4B** and **Table 3S1**).

We use 12 ms as the lower bound for a possible “intermediate latency” chirp response category that includes self-chirps and exogenous small and long chirps. Within the intermediate self-chirp responses, there is a continuum of response latencies between 14.6 and 29.4 ms followed by a large gap to the next latency of 43.8 ms We use a latency at the upper end of this gap, 41 ms, to define an upper bound of the intermediate latency chirp category that is consistent across self- and exogenous small and long chirps (**Fig. 3D, Table 3S1**). There are, within the intermediate category: N=10 self-chirp responses, N=28 small exogenous chirp responses, and N=4 long chirp responses. The intermediate median self-chirp latencies ranged 14.6-29.4 ms (20.6±4.4 ms, mean±SD) while those of exogenous small chirps ranged 12-38.6 ms (24.3±6.9 ms). The means of these distributions are not statistically different (p=0.12, two-sided bootstrap). A latency of 41 ms served as our lower bound for the possible ‘slow response’ category with latencies ranging between 41.8 and 86.2 ms (**Fig. 3D, Table 3S1**). There are, within the slow-response category: N=2 self-chirps, N=3 exogenous small chirps and N=4 long chirp responses.

The same analysis as above reveals that intermediate latency responses for self- and exogenous small and long chirps were typically initiated while the chirp was ongoing or shortly after it ended. In other words, the intermediate latency responses appear to be rapidly evoked by the chirp onset (**Figs. 3, 4** and **Table 3S1**). In contrast, the slow responses were initiated >20 ms after the chirp had ended.

**Figure 4:**
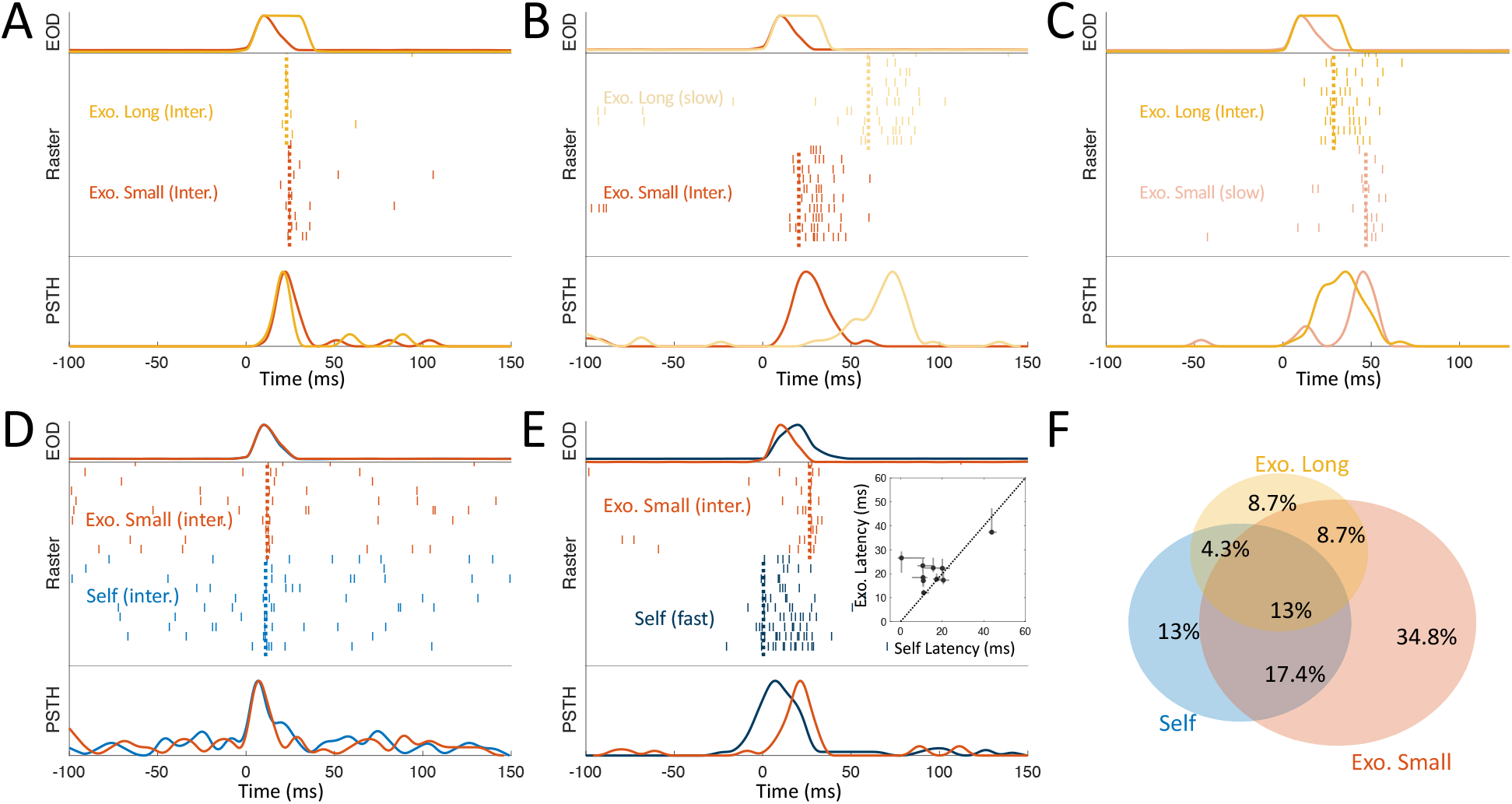
PG cells respond to multiple chirp types. **(A-E)** Top: EOD rate excursion (normalized); Middle: raster plot depicting spike response of one PG cell to individual chirp instances of different kinds (dashed lines-median first spike latency); Bottom: PSTH across chirps. (**A-C**) Three PG cells responding to both small and long exogenous chirps. Cell in A responds with similar latency and pattern, to cells in ELL (Marsat et al., 2009) and TS (Vonderschen & Chacron, 2011); cell in B responds earlier to small chirp, while cell in C responds earlier to long chirp. (**D**) A cell responding to self and exogenous chirp with similar latency, suggesting a common afferent pathway. (**E**) A cell with a fast response to self-chirp (median latency 0.4ms, suggesting motor corollary discharge) and an intermediate response to exogenous small chirp (median latency 26.6ms). Inset: most PG cells responding to both self- and exogenous small chirps responded faster to the former. (**F**) Venn diagram of response to chirp types across the PG population; 56.5% of the cells responded to one chirp only with the remaining 43.5 cells responding to at least two different chirp types.

It is clear that our initial predictions are not met. The fast responses appear to be unique for self-chirps and are initiated before or soon after chirp onset. Further, the first-spike latencies to small chirps lie within a very broad range (median latencies 0.4-65 ms) and not in the narrow range expected from responses in ELL and TS. Responses to long chirps are exceptionally slow (median latency 22.9-86.2 ms). The latency data alone strongly suggest that PG cell responses to the various chirp types are not solely driven by the ascending pathways described in **Fig. 2A**, and do not simply encode the precise chirp time. Moreover, while TS cells respond to small and long chirps with short bursts mostly coextensive with the duration of the chirp (Vonderschen & Chacron, 2011), PG chirp response patterns were highly varied; while some spiking responses faithfully follow their TS input (**Fig. 4A,D**), other responses are clearly different and may persist beyond the chirp duration (**Fig. 4B,C,E**). While we were unable to find any relationship between the response patterns and our proposed divisions based on latency, slower responses exhibited greater chirp-to-chirp variability, a hallmark of longer, multi-synaptic pathways (**Fig. 4S1 A**).

So far, we have described individual response of PG cells to different chirp types. Next, we assess the overlap between these different responses across the PG population. For this purpose, we used cells that had been stimulated with all chirp types – self-chirps and exogenous small and long chirps – to determine how selective these cells are to chirp type (the full dataset upon which the conclusions below are based is presented in **Table 3S1**).

Some PG cells responded to different chirp types with very similar intermediate spike latencies and burst-like patterns (**Fig. 4A, D**, see also **Fig. 4S1 B**), akin to TS responses previously reported (Vonderschen & Chacron, 2011), and we therefore hypothesize that they are simply inherited from the TS cells. However, in other cases cells responded very differently to small vs. long chirps (**Fig. 4B,C**) or to self-vs. exogenous chirps (**Fig. 4E)**. Overall, most of the cells excited by both self- and exogenous-small chirps responded slightly earlier to the former (**Fig. 4E, inset**), suggesting that in these cells the responses to the two chirp types propagated along different pathways (see also examples of complex interactions between PG responses to different chirp types, **Fig. 4S1**). Altogether, about half of PG neurons (56.5%) responded selectively to only one chirp-type, while the other half (43.4%) responded to two or more chirp types (**Fig. 4F**).

Taken together, it is clear that PG is not merely a relay transmitting chirp-related information from TS to pallium but instead massively transforms this information. There are multiple pathways and interactive processing of self- and exogenous small chirps and long chirps. Importantly, the fast self-chirp response category strongly suggests the contribution of a corollary discharge pathway from pre-motor areas involved in chirp generation (see Discussion).

#### Amplitude modulation signals cannot distinguish between Self and Exogenous small chirps

What could be the role of corollary discharge driven responses of the kind reported above? Both self-generated and exogenous chirps create sensory signals across the fish’s skin. To investigate whether fish are able to distinguish between self and exogenous chirps solely based on their sensory signature, we used a numerical model to simulate the electric potential of two interacting fish. The geometry of the fish and their electric organs, as well as their EOD current density profile were taken from a data-calibrated model (Babineau et al., 2006). Gaussian small chirps were generated in the EOD of each of the simulated fish. We quantified the effect of these chirps by measuring the phase shift or ‘reset’ (Benda et al., 2005) they induced in the beat – the periodic pattern of signal amplitude modulations, which is the primary feature driving almost all electroreceptor activity (see **Fig. 5S1** for details). The beat pattern was extracted at various points along the skin of one of the fish (**Fig. 5A**), and the phase reset due to the chirps was evaluated at every position (**Fig. 5B**); a sensory ambiguity would exist if two different social interaction scenarios were to yield the same phase reset curves. We evaluate this possibility by comparing pairs of phase-reset curves associated with self and exogenous chirps for a range of different ΔFs (**Fig. 5C**). Based on this analysis, we conclude that for each self-generated chirp, there exist at least one different interaction scenario where an exogenous chirp yields the same phase reset curve (dark blue regions in **Fig. 5C**). Therefore, the sensory spatiotemporal pattern of amplitude modulations due to a chirp cannot, in and by itself, distinguish between self- and exogenous chirps. This sensory ambiguity may be addressed by complementing the sensory signals with motor-related corollary discharge of the type reported above. We note that while the overwhelming majority of electroreceptors are the so-called P-type receptors that encode amplitude modulations (Carr et al., 1982), a small number of temporal encoders (T-type receptors) (Scheich, 1973), located almost exclusively on the head (Maler, 1979) may respond differently to self- and exogenous chirps. The T-receptor system is, however, poorly developed in Apteronotus sp. (Carr et al., 1982) and may therefore be insufficient to disambiguate self-from exogenous chirps.

**Figure 5:**
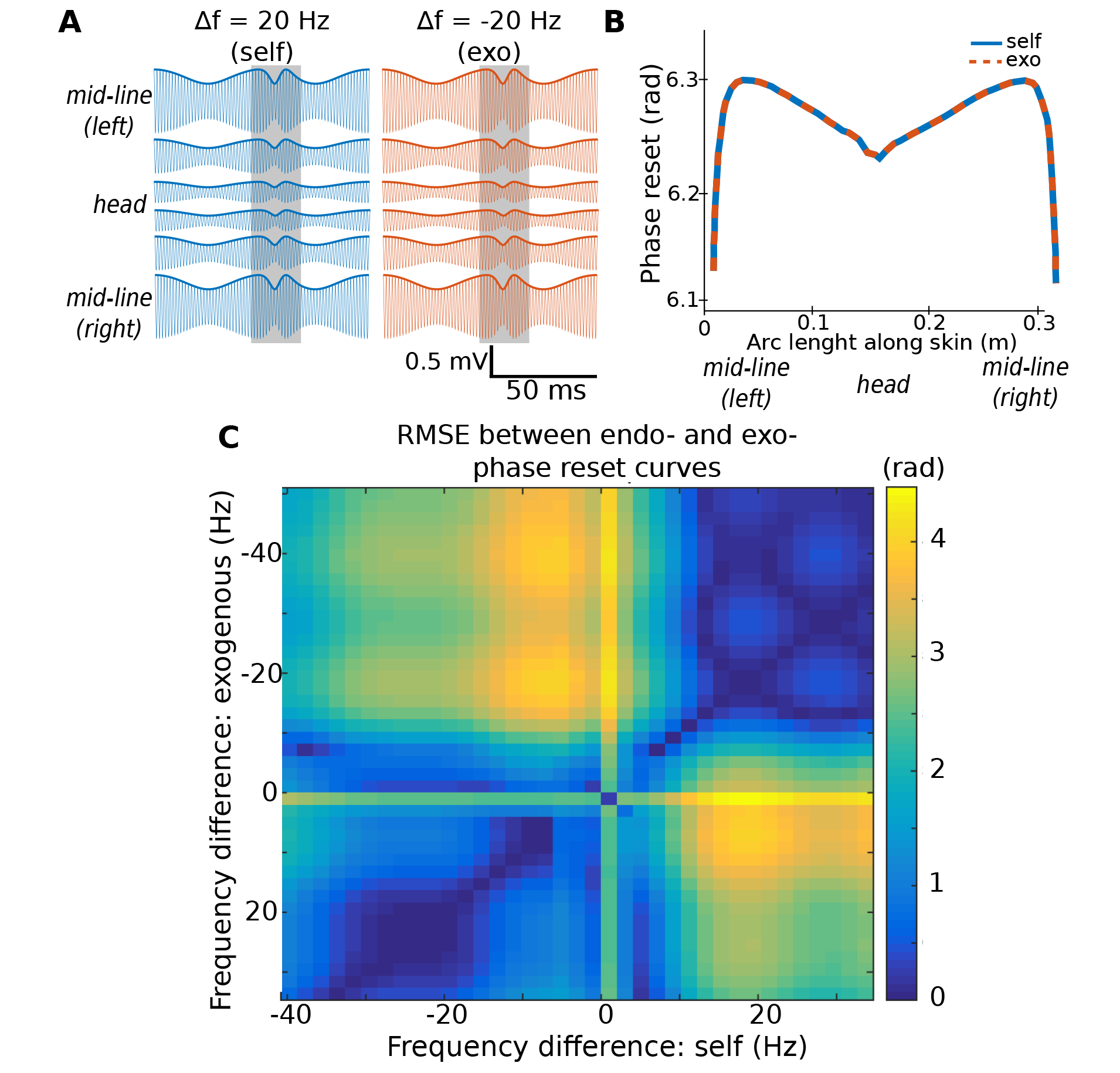
Sensory ambiguity in chirp source identity. (**A**) Simulation of the beat patterns due to the interaction of two fish, measured at different locations along the skin of one of the fish (*fEOD*= 700 Hz). The beat patterns for a self-chirp when *Δf* = 20 Hz (left) is the same as that for an exogenous chirp when *Δf* = −20 Hz (right). The shaded areas indicate when the chirp occurs (peak frequency excursion 80 Hz, duration 15 ms). (**B**) Phase reset due to chirp across the skin surface for self (blue) and exogenous (red) chirp, for the parameters used in (A). Phase reset curves are identical and therefore the two scenarios are indistinguishable using this sensory feature. (**C**) Comparison between phase reset curves for self- and exogenous chirp scenarios. The distance between pairs of curves is defined as the root-mean-square error (RMSE; color coded). Dark blue regions correspond to scenarios where there is little to no difference in phase reset curve between self- and exogenous chirps, leading to a sensory ambiguity as to which fish produced the chirp.

#### PG cells exhibit mixed selectivity to conspecific identity and relative motion

As detailed in the Introduction, conspecific identity can be inferred from the beat pattern generated by the interaction of the two EOD signals (see **Fig. 1**), while the distance to the conspecific and its relative motion can be derived from the beat’s low-frequency envelope signal. How are these two variables – conspecific identity and relative motion – represented in PG? We used a beat-scan protocol to probe PG cell responses to a large range of beat frequencies (i.e., ΔF, the difference between the stimulus frequency and the fish’s EOD frequency) and superimposed envelopes (**Fig. 6S1**). We found numerous PG cells (N=50) that combined coding for the two variables (**Figs. 6 and 7**). In what follows, we first describe the coding of each variable separately and then discuss the joint encoding of both variables.

**Figure 6:**
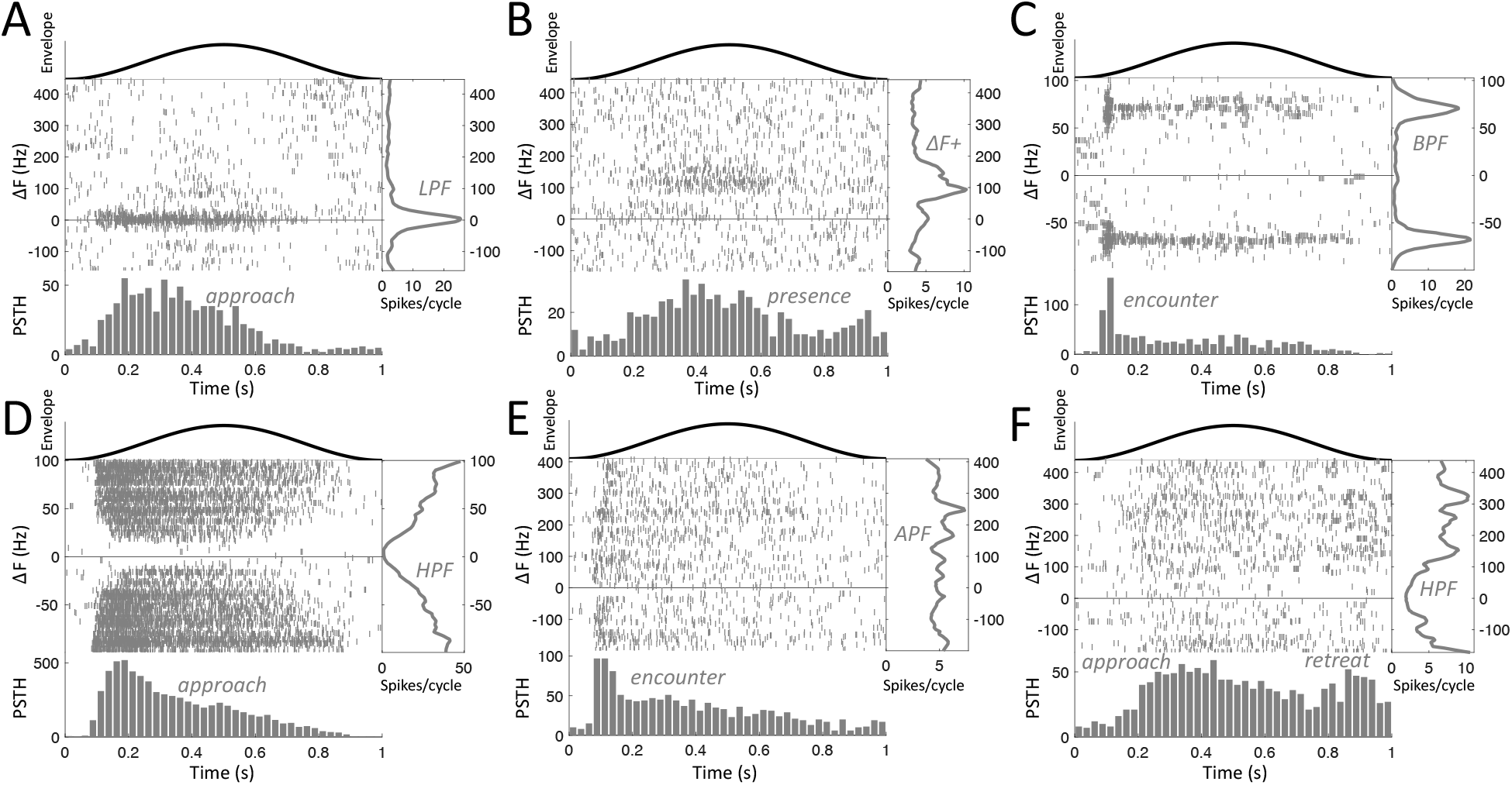
Conjunctive response of individual PG cells to beat frequency and envelope. All panels– middle: raster plot depicting spiking response to each cycle, ordered by beat frequency; top: cycle of the beat envelope waveform simulating approach and retreat; right: number of spikes per cycle vs. beat frequency (i.e., tuning of cell to conspecific identity); bottom: PSTH within the pass band of the cell (i.e., tuning of cell to conspecific relative motion). The different panels show cells responding to: approaching low-frequency beats (**A**), presence of positive frequency beats (**B**), an encounter (envelope onset) within a specific frequency band (**C**), approaching high-frequency beats (**D**), encounter of any beat frequency (**E**), and both approach and retreat of high-frequency beats (**F**). Thus, each cell type has a distinct mixed-selectivity to identity and motion.

**Figure 7:**
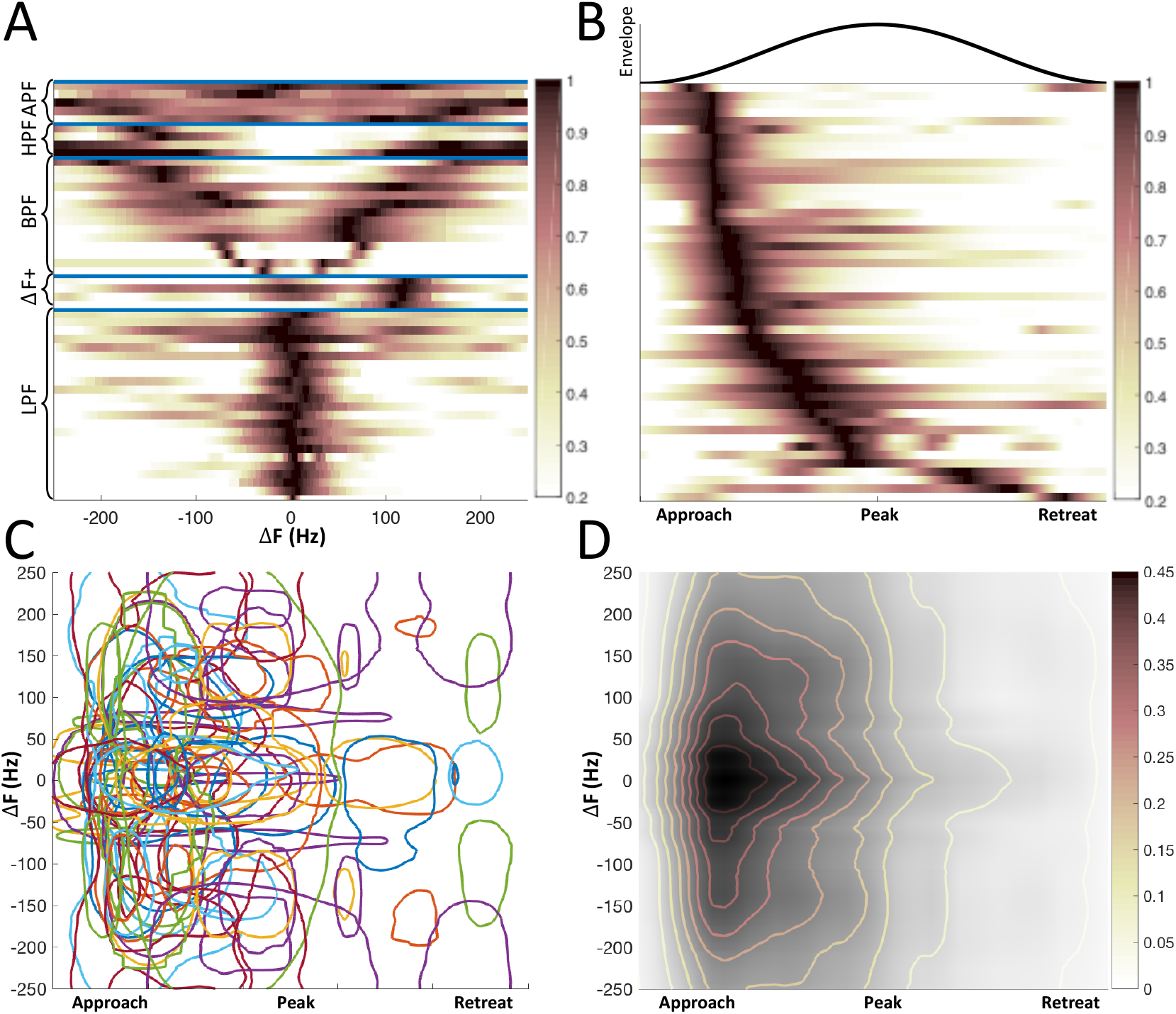
PG population tiles the joint beat-envelope space but overrepresents approaching same-sex conspecifics. (**A**) Normalized tuning to frequency of all beat responding cells, ordered according to response type (from bottom: low-pass, positive frequency, band-pass, high-pass and all-pass filters). (**B**) Normalized PSTH to envelope cycle of all responding cells, ordered according to response phase in the cycle. (**C**) Estimation of the conjunctive response in beat-envelope space; each contour depicts the locus in this two-dimensional space where the cell’s activity is half of its maximal response (band- and high-pass cells have two symmetric contours each). (**D**) Estimation of overall population response.

Each beat-responding PG cell exhibited a preferred range of beat frequencies, and we have identified five frequency tuning categories. First, many cells (N=23) were selective to low frequencies, responding equally to negative and positive ΔF (i.e., representing the absolute frequency difference regardless of sign); these cells therefore acted as *low-pass filters* (LPF, **Fig. 6A**). Second, a few cells (N=4) responded in a narrow frequency range but only at positive ΔFs (i.e., only when the stimulus frequency was greater than the fish’s own EOD frequency (denoted ΔF+, **Fig. 6B)**; engineers call such a response *single side-band filtering*. A third subset of cells (N=14) responded symmetrically within some band around a preferred peak frequency greater than zero, and therefore acted as *band-pass filters* (BPF, **Fig. 6C**). Fourth, a few cells (N=4) were selective to high-frequency beats (*high-pass filters* or HPF, **Fig. 6D and F**). Finally, N=5 cells responded throughout the tested frequency range, albeit not always uniformly, and were therefore denoted *all-pass filters* (APF, **Fig. 6E**). Overall, the PG cell population covers the entire behaviorally relevant beat frequency range, and the ‘filter bank’ provided by the differential tuning across the entire PG population (**Fig. 7A**) can provide the pallium with the necessary information to decode conspecific sex, dominance and identity.

Each of the PG cells also exhibited an envelope tuning, with each firing at a preferred phase of the envelope signal. For each cell, we computed the peri-stimulus time histogram (PSTH, **Fig. 6**, bottom of all panels), averaged across all envelope cycles within the cell’s preferred frequency band. **Fig. 6** included cells responding on the rising phase of the envelope, i.e. as the distance between the fish decreases (*approach* responses, **Fig. 6A, D and F**); cells whose response dynamics track that of the envelope, therefore conveying the distance to the conspecific (reporting its *presence*, **Fig. 6B**); cells strongly and briefly responding at the very onset of the cycle, therefore signaling a new conspecific *encounter* (**Fig. 6C and E**); and finally, a few cells also responded at the declining phase of the envelope (*retreat* responses, **Fig. 6F**). Overall, all phases of interaction are represented in the PG population (**Fig. 7B**), providing the pallium with information of relative conspecific motion. Importantly, while preference of approach phases has been reported to be the predominant feature of brainstem neurons (Huang & Chacron, 2016), brief, encounter-detection responses (which occur at earlier phases of the cycle) were the most common in PG (**Fig. 7B**).

We have shown so far that the same population of PG cells simultaneously encode conspecific identity and relative motion. How is this two-dimensional stimulus space covered? **Fig. 7C** depicts the ‘response region’ of each cell, suggesting that the PG population ‘tiles’ this space, covering all possible combinations of identity and motion. However, it is evident that the coverage is not uniform, with low frequencies and increasing envelope heavily over-represented. This is further demonstrated by estimating the normalized PG population tuning to conspecifics (**Fig. 7D**). This over-representation suggests that PG is especially geared towards detecting and identifying incoming same-sex conspecifics.

#### PG cells exhibit mixed selectivity to chirp signaling and conspecific identity and motion

We have shown how different PG cells convey information about the ‘communication channel’ between fish (the beat pattern and its envelope) and the ‘tokens’ exchanged across this channel (i.e. the various chirp types). What are the interactions between the representation of these different variables? To probe this, we repeated our beat-scan protocol (see **Fig. 6S1**) on a subset of cells (N=22), this time with an embedded chirp (either small or long) in each envelope cycle. Given the limits on recording time, we could only place a chirp at the envelope peak, and so the dependence of the chirp response on relative motion was not probed. This method, therefore, revealed mixed coding for chirps and beat frequency at the envelope peak.

**Fig. 8A** illustrates the responses of a neuron with (purple) and without (green) a chirp delivered at the envelope peak, i.e., when the fish would be in closest proximity. While the cell’s beat response is maximal around ΔF=0 (i.e. this is an LPF cell), its chirp response appears to be confined to a narrow beat frequency range centered around −80 Hz; since the small chirp maximal frequency excursion is 80 Hz, a chirp received on a −80 Hz beat background will bring the beat near 0 Hz, i.e., at the peak beat tuning for this cell. In other words, the apparent ΔF dependence of the chirp appears to result, at least in part, from the chirp frequency excursion bringing the stimulus into the cell’s best beat frequency tuning range. Such interactions between beat and chirp coding have been previously reported in electroreceptors (Walz et al., 2014), and therefore could be inherited by PG. However, not all cells displayed such interactions, with some cells selectively responding only to beats (**Fig. 8B**) or only to chirps (**Fig. 8C**), suggesting further processing in the pathways leading up to PG (Vonderschen & Chacron, 2011). Overall, half of all electrocommunication PG cells responded to both beats and chirps, 20.2% responded only to chirps and the remaining 29.8% only to beats (**Fig. 8D**). Interestingly, the amount of overlap depended on chirp type: while 78% of cells responding to exogenous chirps also responded to beats, only 55% of self-chirp responders responded to beats, providing further evidence that distinct pathways contribute to the responses to the different chirp types. We conclude that PG neurons display mixed selectivity to all components of electric social behavior: conspecific identity, relative distance and motion and incoming and outgoing chirp signals.

**Figure 8:**
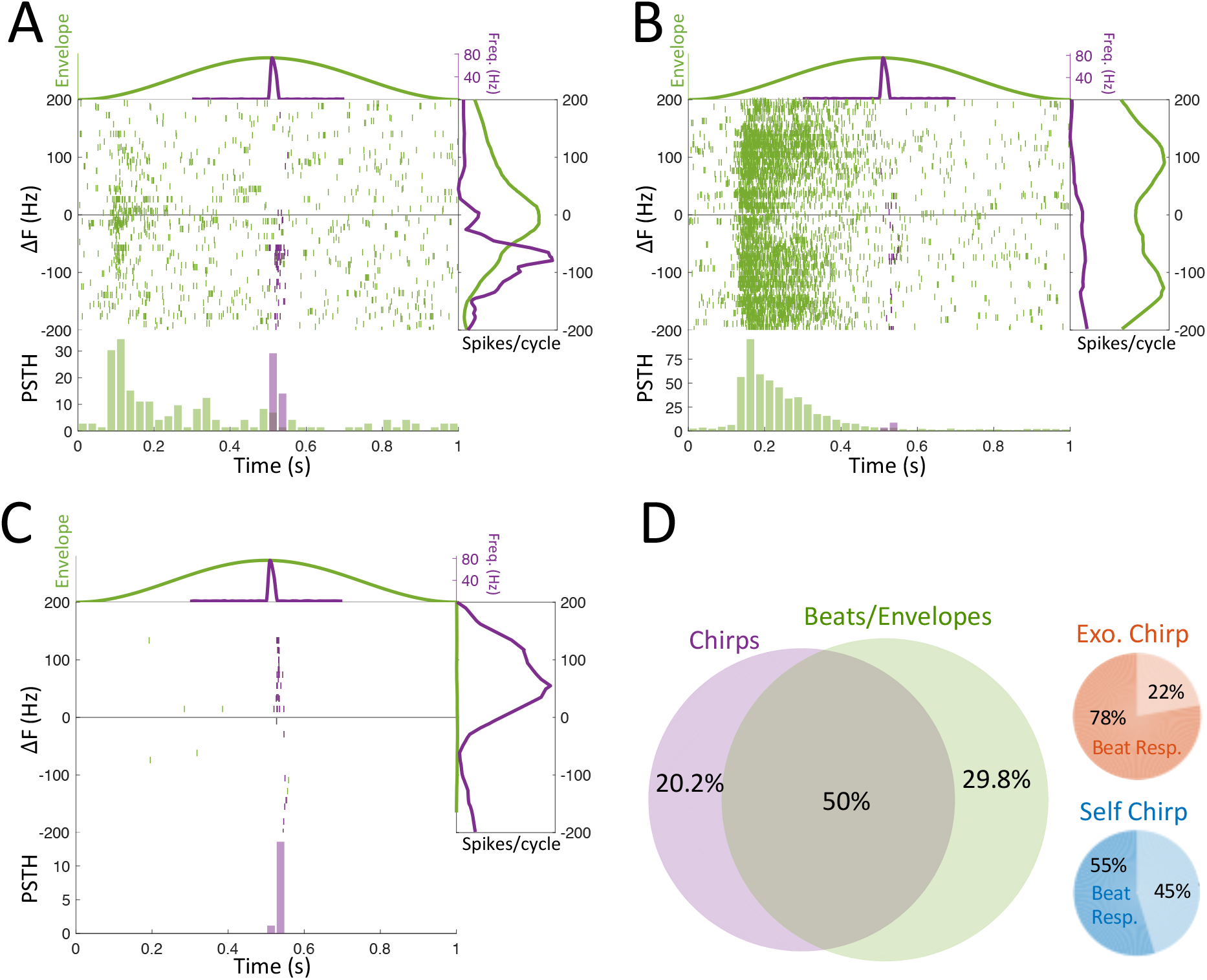
Mixed selectivity to beats/envelopes and chirps. (**A-C**) The beat-scan protocol (see Figure 6S1) was repeated twice, once normally (green) and once with a small/long chirp delivered at the peak of the cycle (purple). middle: raster plot depicting spiking response to each cycle, ordered by beat frequency; top: envelope cycle (left ordinate, green) and frequency excursion (right ordinate, purple); right: normalized spike count per cycle for just the beat (green) and for the chirp (purple); bottom: PSTH. (**A)** A cell whose chirp frequency sensitivity resembles its beat frequency sensitivity shifted by the frequency excursion of the chirp (80Hz). (**B)** A cell responding only to the beat. (**C)** A cell responding only to the chirp. (**D**) Half the PG population responding to social signals responded to both beats/envelopes and to chirps; of the cells responding to exogenous chirp 78% also responded to beats, whereas of those responding to self-chirps only 55% responded to beats as well.

## Discussion

We have demonstrated that the responses of many neurons in the preglomerular nucleus to electrocommunication signals are not simply derived from those of TS, PG’s main source of electrocommunication input. PG neurons respond in a complex manner to combinations of features (beats, envelopes, self and exogenous small and long chirps and corollary discharge linked to small self-chirp emission), with a wide range of response latencies and patterns - we hypothesize that this constitutes mixed selectivity coding for social encounters of Apteronotid fish. Complex mixed selectivity coding has been investigated at higher levels of cortex engaged in ‘complex cognitive tasks’ (Rigotti et al., 2013), and is putatively enabled by strong recurrent connectivity that generates a high dimensional representation of complex tasks (Rigotti et al., 2010). The sensory stimuli associated with primate complex tasks are typically hard to define and the final natural behavioral readout of their representation difficult to quantify. In gymnotiform fish, the stimuli associated with social behavior and electrocommunication can be precisely defined in laboratory and natural habitat settings. The “read out” of the mixed selectivity PG encoding of such signals is, we argue below, carried out by the recurrent pallial networks and results in readily observed “decisions” subsequent to courtship and intruder repulsion behaviors.

Below we first summarize our evidence for the similarities and differences between TS and PG responses to electrocommunication signals and our evidence for a complex mixed selectivity code. We then summarize studies suggesting that chirping behavior is not driven by a direct projection from TS to the prepacemaker (PPn-C) via the nucleus electrosensorius, but rather involves the pallium processing the electrocommunication scene represented by PG activity and controlling chirp initiation via descending projections to the lateral hypothalamus (LH) and preoptic area (PO, see **Fig. 2B,C**). We suggest that LH projections to PPn-C drives small chirps and also provides PG with a corollary discharge signal. Finally, we turn to a study of courtship in the wild of a closely related Apteronotid species (Henninger et al., 2018). We directly infer the electrosensory stimuli generated as a female/male dyad engages in courtship and when the courting male drives off an intruder. We use our inferred stimuli to predict the responses of PG neurons during all phases of these social interactions and the resulting input to pallium. This leads to a direct prediction as to how the activity of the recurrent pallial networks might lead to an apparently “simple” decision on mate choice.

### Combinatorial coding of beats and envelopes

It is useful in the discussion below to distinguish between the signaling properties of beat/envelopes versus chirps. Beats are an obligatory consequence of the difference frequency between the EODs of interacting fish set by their hormonal status. As described above, EOD frequency signals the sex of a mature Apteronotus, and high frequencies are an advertisement for a dominant male (Fugere et al., 2011; Zakon et al., 2002). Electroreceptors can respond to beat frequencies ranging from less than 5 Hz ((Chacron et al., 2005) to at least 425 Hz (Savard et al., 2011) and thereby cover the full range of beat frequencies expected during social interactions. Neurons in the ELL and TS are tuned to specific frequency ranges and respond strongly up to the testing limits used: ELL 120 Hz (Krahe et al., 2008) and TS 256 Hz (Vonderschen & Chacron, 2011). PG neurons are tuned to beat frequencies up to at least 250 Hz (**Fig. 7A**) and a few neurons respond to beats up to 400 Hz (**Fig. 6E,F**). We conclude that PG neurons inherit beat frequency tuning from ascending TS input and can convey information on a conspecific’s sex and status to pallium (**Fig. 2A**).

Envelopes are an obligatory consequence of the relative movement of two or more fish and cannot be set independently of such motion. Electroreceptors can respond directly to envelopes (Savard et al., 2011), and network mechanisms in ELL also generate responses to envelope signals (Middleton et al., 2006). McGillivray et al (McGillivray et al., 2012) showed that a subset of TS neurons responds to envelope signals, but did not specifically address coding for envelope phase. We conclude that PG cells inherit envelope responsiveness from TS neurons, but that a more precise delineation of the transformation of TS to PG responses requires further study.

McGilliveray et al found that different TS neurons were beat frequency tuned, envelope tuned or responsive to both beats and envelopes. In our recordings, the PG beat frequency tuned neurons also discharged at specific envelope phases and vice versa. However, from the joint beat/envelope responses (**Fig. 7C**) it is clear that, for any narrow beat frequency band (e.g., low pass to ±50 Hz), all envelope phases – and therefore motion phases - can be extracted. Likewise, from any narrow envelope-tuned band (e.g., envelope peak tuned), all beat frequencies can be extracted. We conclude that PG jointly encodes beat frequency and envelope so that pallium receives input on the sex, status, and relative motion of an interacting conspecific. Overall, the main transformation of these signals from TS to PG appears to be a transition to a completely combinatorial coding scheme and an over-representation of more ethologically relevant social signals (see below).

### Complex coding of chirps

In contrast to beats and envelopes, chirps are not an obligatory signal – their emission rate is highly variable and depends on hormonal status (Cuddy et al., 2012; Dulka et al., 1995; Dunlap et al., 2002; Zakon et al., 2002), peptidergic innervation (Dulka et al., 1995), monaminergic neuromodulation (Maler & Ellis, 1987; Telgkamp et al., 2007), social context (K. Dunlap & J. Larkins-Ford, 2003; Henninger et al., 2018; Tallarovic & Zakon, 2002), and past experience (Allen & Marsat, 2018; Harvey-Girard et al., 2010). Responses of PG cells to small and long chirps were very diverse with respect to their latency and spiking pattern, suggesting the contribution of at least three separate pathways.

The most straightforward route is the ascending ELL->TS->PG pathway, which generates responses that match these upstream regions (Marsat et al., 2009; Vonderschen & Chacron, 2011), i.e., transient spike bursts with an intermediate median latency whose duration matches that of the chirp. This category of PG neurons would inform pallium of the precise time of occurrence of a chirp, but not of its source (self or exogenous, **Fig. 4D, Fig. 4S1 B**) or even its type (small or long, **Fig. 4A**).

A second pathway underlies the responses to self-chirps that are too fast for transmission via the ELL->TS->PG pathway (median latency: 0.4 to 11.3 ms; minimal latency: −6 ms - +9 ms). We hypothesize that these responses to self-chirps are not due to electrosensory input, but rather to corollary discharge of LH cells that project to both the ventral dendrites of PPn-C cells and to PG. Our arguments for this hypothesis are presented in detail below. We propose that these cells can distinguish self from ‘other’ chirps and, because of their fast latency, modify the processing of exogenous chirps with longer latency responses (**Fig. 9E-G**). Exemplars of this putative category are illustrated in **Figs. 3A and 4E**. Our computational model shows in fact that there exists a sensory ambiguity between self and other chirps, for both positive and negative frequency differences (**Fig. 5**); this arises from the physical nature of the electrical beat interactions and their modulation by chirps. The proposed corollary discharge could thus disambiguate between different chirp sources.

**Figure 9:**
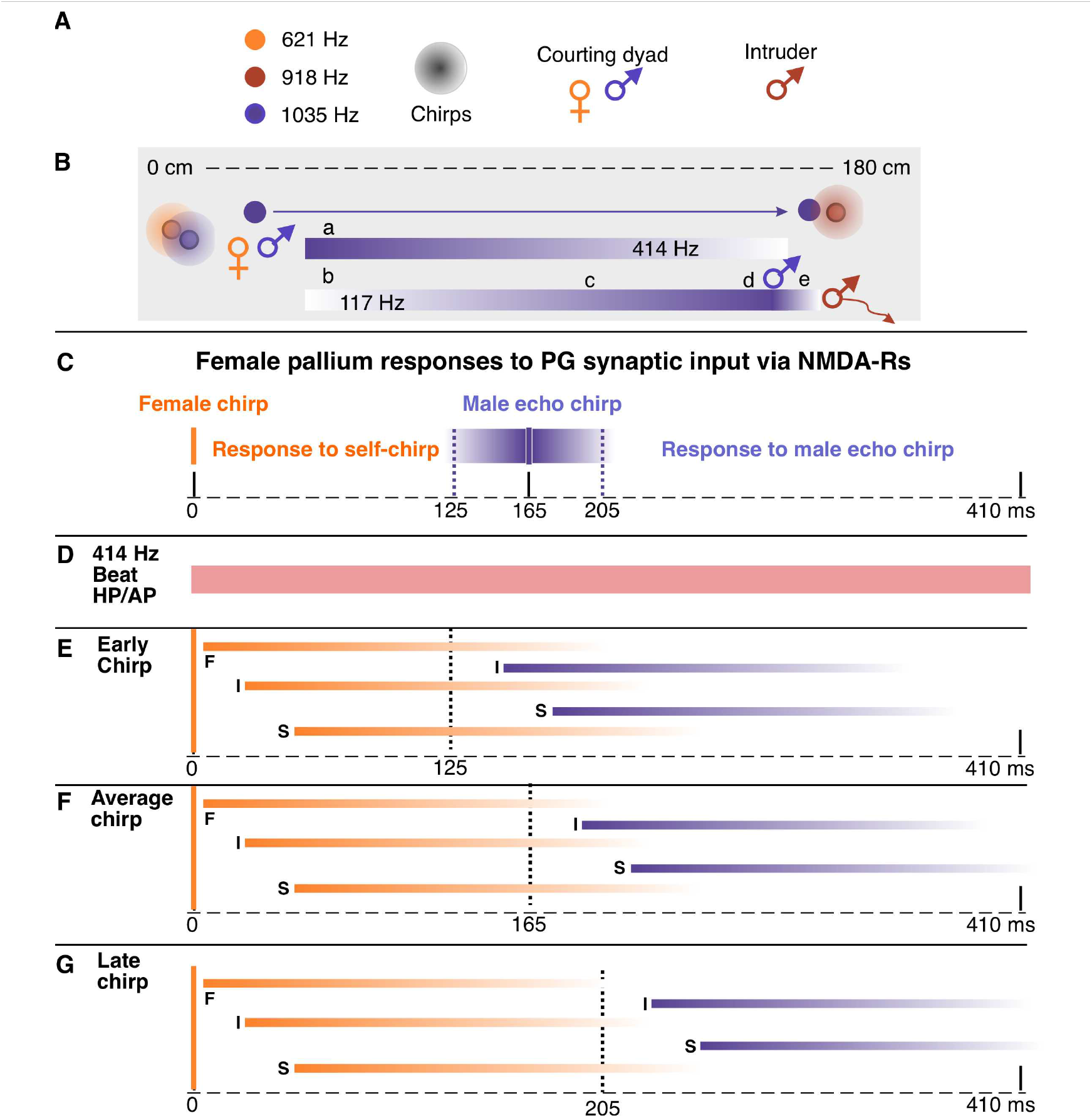
PG representation of natural social interactions. (**A**) The scenario described by Henninger et al and using the graphics of their Figure 2. The depth of the color is proportional to the amplitude of the beat. A courting dyad (male: 1035 Hz; female: 621 Hz) generate a 414 Hz beat frequency. An intruder male (918 Hz) appears during courtship. (**B**) The courting male detects the intruder at a distance of ∼180 cm and swims directly towards him. This generates two envelope signals: one on the 414 Hz female/male beat and another on the 117 Hz male/male beat. As the male leaves the female a retreat envelope (a) is generated on the 414 Hz beat. As the courting male approaches the intruder, encounter (b), approach (c) and presence (d) envelope signals are generated on the 117 Hz beat. The intruder male swims away and emits a few small chirps on the retreat envelope (e). (**C**) The female emits small chirps (vertical orange bar) at a rate of ∼1Hz and the male emits echo small chirps (vertical blue bar) with a mean latency of 165 ms and a standard deviation of 20 ms. The blue shaded bars represent the distribution of echo chirp latencies and the dashed lines two standard deviation latencies: 125 ms and 205 ms. (**D**) The 414 Hz beat of the courting dyad is encoded by high pass (HPF) and all pass (APF) PG cells and transmitted to pallium for the entire duration of courtship. (**E-G**) The PG input to pallium is assumed to utilize NMDA receptors. Activation of these receptors evoked long-lasting EPSPs whose amplitude is reduced to ∼20% of maximum by 200 ms post stimulation (Berman et al., 2001). The time course of these EPSPs is indicated by orange (estimated pallial responses to female self-chirps) and blue (estimated pallial responses to male echo chirps). The predicted synaptic responses with fast (F), intermediate (I) and slow (S) latency responses are shown. (**E**) An early male echo chirp (125 ms) results in extensive overlap of the putative pallial cell responses to female and echo chirps. (**F**) Overlap is reduced for the average chirp latency. (**G**) There is minimal overlap for the late chirp response.

Lastly, we have a heterogenous category of PG chirp responses exhibiting very long latencies (>45 ms) and/or irregular spike patterns. The pathways generating these responses are not known and might include: (a) multiple stage interactions within TS (Carr & Maler, 1985) and/or tectum (Heiligenberg & Rose, 1987; Sas & Maler, 1986) followed by TS/tectum projections to PG; (b) TS/ central posterior nucleus (CP)-> n. electrosensorius (nE)-> nucleus tuberis anterior-> PG (Giassi et al., 2007); (c) descending pallial input to PG. Exemplars of this putative category are illustrated in **Fig. 3 and Fig. 4B,C**. Based on the dispersal of latencies and alterations in spike response patterning compared to TS (Vonderschen & Chacron, 2011), we hypothesize that these pathways perform complex and perhaps nonlinear transformations on the ascending afferent chirp input. The mechanisms of transformation and the potential role of a latency code for electrocommunication signals will be an important focus for future studies.

### Complex mixed selectivity coding of beats/envelopes and chirps

One half of PG neurons in our sample responded to some combination of beat/envelope and chirp signals, with the remaining cells responsive to beat/envelope (∼30%) or chirps (∼20%) alone. Our sample set is too small for refined statistical analyses of the various possible beat/envelope + chirp categories. However, inspection of **Table 3S1** demonstrates that beat/envelope responsive cells can also respond to small (self and exogenous) and long chirps and that they can do so over a wide range of chirp response latencies. Our results are consistent with a random mixing of responsiveness across chirp categories and beat/envelope signals. This hypothesis further implies that parts of the pallium (dorsolateral, DL and/or dorsomedial, DM) encode electrocommunication signals in a high dimensional coding space. The hypothesized purpose of a high dimensional coding space is to facilitate read-out of complex signal combinations (Johnston et al., 2020). ‘Read-out’ in this case implies that the pallial networks control both locomotor and chirping responses to electrocommunication signals. Below we provide evidence that beats/envelopes and chirps do not reflexively cause chirp responses, but rather act via pallium and diencephalic cell groups, i.e., ‘interpreting’ beats/envelopes and chirp afferent inputs and using this ‘interpretation’ to guide chirp and motor outputs is a complex cognitive operation.

### Electrosensory input does not directly evoke chirping

Early studies suggested that there was a one-way link between electrosensory input and chirp output: TS projects to nE and it, in turn, projects to CP and PPn (Carr et al., 1981; Keller et al., 1990; Walz et al., 2012). This projection was hypothesized to drive gradual (PPn-G) and fast (chirps, PPn-C) increases in EOD frequency (**Fig. 2A**) (Metzner, 1999). Heiligenberg et al (Heiligenberg et al., 1996) demonstrated that, in Apteronotus, applying glutamate to a subnucleus of nE (nE-up, nE↑) consistently and immediately caused a gradual increase in EOD frequency. This is consistent with Kawasaki et al’s (Kawasaki et al., 1988) demonstration that glutamate application in the nE↑ target, PPn-G, also causes such increases. This strongly suggests that the nE↑?to PPn-G pathway is the physiological basis of the slow rises seen during Apteronotus conspecific interactions (Zakon et al., 2002). There is a strong direct projection from dorsal central pallium (DC) that terminates precisely on nE↑ (Giassi, Duarte, et al., 2012), and we further propose that this is a route by which pallium can control production of slow rises (**Fig. 2A**).

Heiligenberg et al (1996) showed that glutamate application to nE↑ also caused occasional and very long latency chirping (>2 s). We suspect that the evoked chirping might be an artefact caused by slow diffusion of glutamate to the lateral dendrites of PPn-C cells (Kawasaki et al., 1988), but further work is required to settle this issue. The following data suggests that evoked chirping behavior is independent of nE↑.

#### Spontaneous chirping

Maler and Ellis (1987) and Engler et al (2000) showed that Apteronotid fish can emit spontaneous small chirps, i.e., without any electrosensory input and therefore without any activity in nE/nE↑ (Heiligenberg et al., 1991; Heiligenberg et al., 1996). Notably, Maler and Ellis (1987) showed that the rate of spontaneous chirping was greatly increased by noradrenaline; we hypothesize that this due to the very strong noradrenergic innervation of LH (Sas et al., 1990) (**Fig. 2B,C**). Notably, Engler et al reported (Engler et al., 2000) that, after evoking chirps with a single short (30 s) stimulus, the rate of spontaneous chirping increased dramatically for several minutes post-stimulus; this effect was specific to chirping and did not increase the rate of slow rises.

#### Long latencies between beat stimulus presentation and evoked chirping

Stimulating a fish with an EOD mimic sine wave causes beats that can evoke chirping. There is a very long latency between the onset of a beat stimulus and evoked chirping in males (1-25 s) (Zupanc & Maler, 1993) and females (91 s mean) (Triefenbach & Zakon, 2003).

#### Long latencies of echo chirp responses

The chirp of one fish can evoke a chirp in a nearby conspecific. Latencies of these ‘echo chirps’ were found to occur randomly between 385-4103 ms (Zupanc et al., 2006) and 200-600 ms (Hupe & Lewis, 2008) for males under laboratory conditions; mean echoing latency of a courting dyad in the wild was 165 ms (SD 20 ms) (Henninger et al., 2018).

The occurrence of spontaneous chirps and the long and variable latencies of beat- and echo-evoked chirps are not consistent with a direct TS-> nE↑-> PPn-C pathway driving chirping. We propose below that chirping – including echo-chirping – is driven by an autonomous network interconnecting subpallium (SP), PO, LH and CP (**Fig. 2**). We further propose that electrosensory input reaches this network via PG projections to pallium. The main inputs to PPn-C (chirp cells) relevant to this paper are SP, PO, nE (especially nE↑), CP and LH (Wong, 1997a, 1997b). Of these regions, SP (Wong, 1997b), CP, LH and PO also project to PG (Giassi, Duarte, et al., 2012), and are therefore possible sources of the fast, putative corollary discharge driven self-chirp responses in PG. The SP nuclei (ventral ventral Vv, ventral dorsall Vd) are composed of GABAergic neurons (Mueller & Guo, 2009) and therefore can be ruled out (see below). CP can be ruled out because glutamatergic stimulation within this region causes only slow rises (Kawasaki et al., 1988). We therefore discuss LH and PO as both chirp generators and possible sources of corollary discharge to PG. We also discuss the role of SP because of its intimate interconnectivity with all these nuclei and a potential link to the behavioral data of Henninger et al. (see below).

### Hypothalamus drives small chirps and provides a corollary discharge input to PG

Glutamate application throughout the extensive ventral dendrite domain of PPn-C neurons induces barrages of small chirps (Kawasaki et al., 1988). The ventral dendrites of PPn-C neurons receive multiple peptidergic inputs (Richards & Maler, 1996; Sas & Maler, 1991; Weld & Maler, 1992; Yamamoto et al., 1992) sweeping up from LH (Wong, 1997b). The expression of Substance P is sexually dimorphic with far greater expression in the territory of the PPn-C ventral dendrites, LH and PO of males (Weld & Maler, 1992). There is strong evidence linking LH Substance P cell projections to PPn-C to chirping (Dulka & Ebling, 1999; Dulka et al., 1995; Weld & Maler, 1992); we do not know the possible role(s) of the other peptidergic inputs in the control of chirping. We hypothesize that neurons in LH are the main, and perhaps only, drivers of small chirps. We hypothesize that the strong LH to PG projections (Giassi, Duarte, et al., 2012) are a corollary discharge pathway that causes the fast responses to self-chirps. We predict that stimulation of LH will cause chirping and evoke spiking in PG neurons. Further, the latencies to evoked chirping and spiking will be similar and short enough to be consistent with monosynaptic projections from LH to PPn-C and PG.

### Vasotocin expressing preoptic neurons may drive long chirps and provide a corollary discharge input to PG

Pouso et al has described vasotocin expressing neurons in the PO of gymnotiform pulse species (Gymnotus omarorum and Brachyhypopomus gauderio) (Pouso et al., 2017). These neurons projected to multiple sites including PPn and PG, confirming earlier findings (Giassi, Duarte, et al., 2012; Wong, 1997a). Pouso et al then used cFos expression to show that PO vasotocin cells were activated by social interaction in a reproductive context (Pouso et al., 2019) in a pulse species (Brachyhypopomus gauderio), and suggested that these neurons were active during courtship. Most importantly, Bastian et al have shown that, in Apteronotus, intraperitoneal injections of vasotocin cause some males to emit long chirps (Bastian et al., 2001). The long chirps were preferentially induced by high frequency beats typical to opposite-sex interactions (Hagedorn & Heiligenberg, 1985). Bastian et al (2001) also report preliminary experiments showing that stimulation of PO will evoke long, but not small, chirps. We hypothesize that the PO projections to PG are a corollary discharge pathway for long chirps. We predict that stimulation of the PO will cause, in a fish in a suitable endocrine state, long chirps and spiking in PG neurons. Further, the latencies to evoked chirping and spiking will be similar and short enough to be consistent with monosynaptic projections from PO to PPn-C and PG.

Finally, we hypothesize that the subpallium inhibits chirping via its projections to LH and CP/PPn-C and an inhibitory corollary discharge to PG. Wong found that electrical stimulation of PO induced chirping, but only after a latency of at least 500 ms (Wong, 2000). Importantly, glutamatergic stimulation at the same site did not have any effect. Following Wong, we interpret this result as implying that stimulation activated fibers of passage from SP nuclei (Vv, Vd) to PPn-C (Wong, 1997a, 1997b). These subpallial nuclei are now known to be GABAergic (Mueller & Guo, 2009) and likely homologous to the basal ganglia and septal region of tetrapods (Elliott et al., 2017; Giassi, Ellis, et al., 2012; Harvey-Girard et al., 2013; Toscano-Marquez et al., 2020). We therefore hypothesize that SP inhibits chirping via its projections to PO (long chirps) and LH (small chirps) and simultaneously inhibits PG cells; the possible consequences of such inhibition for courtship behavior are mentioned below.

We hypothesize that the interconnected network illustrated in **Fig. 2C** – SP, PO, LH and CP/PPn-C – constitutes the machinery that organizes chirp patterning during social interactions, and simultaneously provides positive and negative corollary discharge to PG. These nuclei do not receive direct input from PG. Below we review evidence that pallium (DL and/or DM) intervenes between PG responses to beat/envelopes and chirps to control this machinery.

### Descending pallial projections drive electrocommunication signaling and modulate electrocommunication afferent pathways

The vast majority of PG neurons respond to electrosensory stimuli associated with electrolocation (Wallach et al., 2018) or electrocommunication (this paper). These neurons project massively to both DL and DM. The descending projections of the pallium include those that might control chirping (LH, PO, **Fig. 2B**), slow rises (nE**↑ Fig. 2A**) and electrocommunication afferent input to pallium (TS, PG, **Fig. 2A,B**). There is direct evidence for DL involvement with electrocommunication (see below), but published evidence is lacking for DM. A recent study in a pulse gymnotiform fish (*Brachyhypopomus gauderio*) has shown that social interactions, including electric signaling, induces strong expression of an immediate early gene in DM neurons (P. Pouso and A. Silva, personal communication, January, 2021). This provides direct evidence for DM involvement in social behavior. Given the lack of information on intrinsic DM circuitry and physiology, we do not further discuss its potential role(s).

There is already direct evidence, in addition to the circuitry illustrated in **Fig. 2B**, that DL processes the electrocommunication signals that it receives from PG and that its descending output can adaptively control both electrocommunication processing and output. Santana et al reported that blocking GABAergic transmission in DL caused large complex variations of EOD frequency in a pulse gymnotiform fish (*Gymotus carapo*) (Santana et al., 2001). Harvey-Girard et al reported that stimulating *Apteronotus* with an EOD mimic evoked chirping and caused strong expression of an immediate early gene (Egr-1) in DL, DD and DC (Harvey-Girard et al., 2010). The fish’s chirping to a particular ΔF habituated and, in parallel, Egr-1 expression was reduced. Stimulation with a different ΔF or at a different location re-induced Egr-1 expression. Elliott and Maler recorded from DD of Apteronotus and reported that stimulation with a sine wave EOD mimic induced Up states and spiking discharge in DD neurons demonstrating functional consequences of the connectivity illustrated in **Fig. 2B** (Elliott & Maler, 2015).

The critical paper is from Huang et al and follows a series of papers on ELL responses to envelope signals in Apteronotus (Huang et al., 2019). Huang and Chacron showed that ELL pyramidal cells could optimally encode envelope signals (Huang & Chacron, 2016). Huang et al and Metzen et al then showed that optimization required feedback input from a rhombencephalic nucleus (Huang et al., 2018; Metzen et al., 2018). Envelope statistics depend on the motions of two or more interacting fish (Fotowat et al., 2013; Yu et al., 2012). Huang and Chacron (2019) then reported that ELL pyramidal cells could adapt to changes in envelope statistics so as to maintain optimized coding; adaptation took over 60 minutes. Most importantly, lesions of the entire telencephalon prevented pyramidal cell adaptation to the new envelope statistics; pyramidal cells continued to respond to the initial envelope statistics and therefore no longer optimally encoded the new stimulus. Based on the known circuitry and physiology (**Fig.2A**, Huang et al 2018; Metzen et al 2018), Huang et al (2019) logically proposed that telencephalic control of adaptation is via descending projections to TS. Our data strongly suggests that envelope signals encoded in PG reach DL and consequently DC; DC projects to the TS (Giassi, Duarte, et al., 2012) which can in turn access the rhombencephalic nucleus required for optimization of envelope coding (Carr et al., 1981; Clarke & Maler, 2017). We predict that a subset of DC neurons, likely those expressing AptFoxP2 (Harvey-Girard et al., 2012), will respond to envelope signals and that selective lesions of DC will prevent adaptation of ELL pyramidal cells to changes in envelope statistics.

We conclude that the link between electrcommunication signal input and electrocommunication output (chirping and locomotor) does indeed run from PG to pallium and then to LH, PO and SP to control small and long chirp production. The pallial descending projections to TS and PG can also control pallial input. **Fig. 2** summarizes these connections.

### High dimensional coding of electrocommunication signals: implications for behavior

We finally turn to a detailed seminal study of natural social behavior of *Apteronotus rostratus*, a member of the *Apteronotus leptorhynchus* species group, in the wild (Henninger et al., 2018), in order to highlight potential social computations enabled by the mixed selectivity coding in PG. Henninger et al report that (see their Fig. 4) “Roaming males approached and extensively courted females by emitting large numbers of small chirps” as well as agonistic interactions between pairs of males and pairs of females. Females can be courted by several males per night and, while this was not explicitly reported, we assume that not all males are accepted by a female.

An exemplary scenario described in Henninger et al (see their Fig. 2) is reproduced in **Fig. 9**. A male (EOD: 1035 Hz) and female (EOD: 621 Hz) are courting in close proximity (beat: 414 Hz) (**Fig. 9A**, “a”). An intruder male (EOD: 918 Hz) interrupts courtship at a distance of ∼180 cm (**Fig. 9B**). We hypothesize that the onset of an envelope (“b”) of the 117 Hz beat frequency first alerts the courting male; the over-representation of this ‘intruder encounter’ scenario in PG suggests its ethological importance. The courting male then swims directly towards the intruder and experiences a receding envelope on the 414 Hz female beat and an approach envelope on the 117 Hz male beat (“c”). We note that, while the stimuli are related to the social encounter, the control of swimming is likely from a subset of AptOtx1 expressing DC neurons projecting to the tectum (Giassi, Duarte, et al., 2012; Harvey-Girard et al., 2012). Tectal projections to the reticular formation (Heiligenberg & Rose, 1987) and reticular formation projections to spinal cord (Behrend & Donicht, 1990) presumably direct the courting male towards the intruder – electrocommunication and electrolocation computations are linked at the level of the pallium and tectum.

The envelope reaches its peak when the courting male reaches the intruder male (“d”); we suggest that the fast approach envelope of the 117 Hz beat (no chirps) constitutes an aggressive signal. The intruder male is then driven away, emitting small chirps on a retreat envelope (“e”); Henninger et al interpret this as a submissive signal that deters aggression, see also (Hupe & Lewis, 2008). We hypothesize that this behavioral sequence – small chirps on the background of a beat with frequency typical to same-sex interactions and a decreasing envelope – is represented as a three-dimensional trajectory (beat+envelope+chirps) in a high dimensional coding space within the male pallium, signifying a submissive signal.

The courting male then returns and resumes courting the female **(Fig. 9C**). The courtship sequence involves the female emitting small chirps in response to the beat but independent of the male’s chirps. The male echoes the female’s small chirps at an average latency of 165 ms (SD 20 ms). From the high dimensional coding space perspective, the female’s pallium interprets the PG activity due to the combination of the 414 Hz beat and her own (corollary discharge driven) self-chirp input to assign the male echo chirps to a “courtship” category rather than a “submission” or “aggression” category. The female then emits a single long chirp and chirping diminishes or ceases for several seconds before it re-commences; inhibitory input from SP might cause the reduction in chirping (**Fig. 2C**). Courtship consists of multiple such female chirp/male echo bouts.

We assume that the female is using the male echo response to select mating partners. There are three simple possibilities for how the female evaluates a potential mate: the probability, the latency or the temporal precision of the male’s echo chirp response. The critical point is that the Apteronotus pallial neurons strongly express NMDA receptors (Bottai et al., 1997; Harvey-Girard et al., 2007; Maler & Monaghan, 1991). Activation of NMDA-Rs of ELL pyramidal cells evokes an EPSP lasting approximately 200 ms (Berman et al., 2001). As illustrated in **Fig. 9E-G**, PG cell responses to a female’s self-chirps should cause strong activation of their DL target network for >125 ms. The response of DL neurons to the male’s echo chirp (**Fig. 9E-G**) will, we hypothesize, depend on the latency of the male’s echo chirp – the shorter the echo chirp latency, the stronger the summed evoked responses. This is a simple consequence of the long duration of NMDA-R mediated EPSPs permitting summation of the responses to the female’s self-chirp and the echo chirp at short echo latencies (**Fig. 9E**); a missing echo chirp would therefore lead to the weakest activity of the post-echo chirp DL activity. We hypothesize that a female will consider a high probability of consistent and precisely timed short latency echo chirps as indicating a fit male. We note that the mixed selectivity coding of PG, reflected by the extensive overlap of self and exogenous chirp responses (**Table 3S1** and **Fig. 4**), as well as chirp and beat/envelope responses (**Fig. 8**), suggests that the DL recurrent network is able to selectively detect the combined scene of a high frequency beat, a self-chirp, and an echo chirp with a particular latency.

Temporal and spatial propagation of DL-intrinsic activity will depend on the local recurrent architecture of DL (**Fig. 2B**) (Giassi, Ellis, et al., 2012; Trinh et al., 2016) and the intrinsic dynamics of DL neurons (Trinh et al., 2019). For example, long-lasting self-chirp induced DL reverberatory activity ((i.e., >200 ms), (Trinh et al., 2019; Trinh et al., 2016) might greatly amplify overlapping female + echo chirp responses. A similar scenario might apply to the effect of the reciprocal connections between DL and the globally recurrent DD network (**Fig. 2B**) (Elliott et al., 2017; Elliott & Maler, 2015; Giassi, Ellis, et al., 2012). Finally, we propose that pallial circuitry and dynamics represents the courtship sequence as a trajectory in a high dimensional coding space within the female pallium. The culmination of this trajectory converts the electrosensory input into emergent ethological “meaning” for the female. One meaning is “Fit Male” and the associated action is laying an egg and coordinating its fertilization. The other possible meaning is “Unfit Male” and results in rejection.

### Parallels with mammalian circuits for social behavior

We finally propose that our main conclusions are entirely similar to recent ideas concerning mammalian social behavior neural circuitry:

#### The preoptic area

Stimulation within PO evokes sexual behavior in mice (Wei et al., 2018), including the production of male courtship ultrasonic vocalizations (Gao et al., 2019). PO projects to many sites including the paraventricular nucleus of the thalamus (PVN) (Kirouac, 2015; Vertes et al., 2015).

#### The ventromedial hypothalamus

The ventromedial hypothalamus (VMH) governs mating and social aggressive behavior in mice (K. Hashikawa et al., 2017; Y. Hashikawa et al., 2017; Lo et al., 2019; Yang et al., 2013). This cell group also projects to many targets including PVN (Y. Hashikawa et al., 2017; Kirouac, 2015).

#### Descending control of mating/courtship and aggression

PVN has widespread cortical projections including parts of prefrontal cortex and the hippocampal formation (Y. Hashikawa et al., 2017; Vertes et al., 2015), which in turn project back to both PO and VMH (Cenquizca & Swanson, 2006; Kishi et al., 2000; Saper, 2000).

We propose that the following are ancestral vertebrate circuits for control of social communication and behavior:

a. Hypothalamic and preoptic cell groups are the command centers for courtship, mating and aggressive social behaviors in mammals and Apteronotus.
b. These cell groups transmit corollary discharge outputs to thalamus (PVN in rodents, PG in Apteronotus).
c. The pallial/cortical targets of these thalamic cell groups (hippocampus, prefrontal cortex/DL, DM) then send return projections to the preoptic area and hypothalamus. The possible function of such descending projections with respect to aggression is succinctly summarized by Y. Hashikawa et al (2017): “Given that dominance behavior depends critically on evaluating self and opponent, changes in dominance behavior may reflect changes in perceived relationship of an opponent to oneself.” In other words, the pallial/cortical projections to PO and LH/VMH provide the learned contextual information required for vertebrate males to decide who is dominant and for females to decide who is a good mate, and to then take the appropriate actions – fight or retreat in the former and accept or reject in the latter. Weakly electric fish, with their stereotyped and readily recorded and mimicked electrocommunication signals and smaller and less complex brains, are an excellent preparation for studying the neural networks responsible for social behaviors and decisions.

### Thalamic encoding of social signals in electric fish as a framework for studying neural representation of high-dimensional coding spaces

A recent study recorded spontaneous activity from more than 11,000 neurons in the visual cortex of a mouse (Stringer et al., 2019). Sophisticated statistical analyses showed that over 120 dimensions were required to account for approximately 90% of the discharge variability across the population. A key conclusion of this paper is that cortex utilizes a mixed high dimensional code to represent visual images. We speculate that recording across all of cortex while a mouse was engaged in complex natural social behaviors would require a staggeringly large number of dimensions. There is, at present, no way to connect the various coding dimensions to specific combinations of percepts, choices and actions, and there is also no way to understand how the incredibly complex cortical circuit dynamics can generate coding spaces with 100s of dimensions.

In contrast, the limited number of electrocommunication signals and associated locomotor actions suggests that the number of behavioral dimensions comprising courtship and rivalry in electric fish is quite small. In this paper, we have mixed selectivity coding of this multi-dimensional space already at the level of PG, the thalamic bottleneck through which all sensorimotor information is relayed to the pallium, itself a much smaller and simpler structure compared to the mammalian cortex (Elliott et al., 2017; Giassi, Ellis, et al., 2012; Trinh et al., 2016). We therefore suggest that the pallium of weakly electric fish and its involvement in social interactions would be a highly beneficial system to obtain deeper understanding of how neural circuit dynamics can generate complex, possibly nonlinear, high dimensional coding spaces.

## Acknowledgements

This research was supported by an CIHR grant to LM and AL (CIHR# 153143) and NSERC grants to LM (RGPIN/04336-2018) and AL (RGPIN/06204-2014).

## Methods

### Experimental Model

All procedures were approved by the University of Ottawa Animal Care and follow guidelines established by the Society for Neuroscience. *Apteronotus leptorhynchus* fish (imported from natural habitats in South America) were kept at 28°C in community tanks. Fish were deeply anesthetized with 0.2% 3-aminobenzoic ethyl ester (MS-222; Sigma-Aldrich, St. Louis, MO; RRID: SCR_008988) in water just before surgery or tissue preparation.

### Surgical procedure for in-vivo recordings

Surgery was performed to expose the rostral cerebellum, lateral tectum and caudal pallium. Immediately following surgery, fish were immobilized with an injection of the paralytic pancuronium bromide (0.2% weight/volume), which has no effect on the neurogenic discharge of the electric organ that produces the fish’s electric field. The animal was then transferred into a large tank of water (27 °C; electrical conductivity between 100–150 μS cm^−1^) and a custom holder was used to stabilize the head during recordings. All fish were monitored for signs of stress and allowed to acclimatize before commencing stimulation protocols.

### Neurophysiology

Custom made stereotrodes or tritrodes were made of 25-μm diameter Ni-Cr wire (California Fine Wires). Each electrode was manually glued to a pulled filamented glass pipette (P-1000 Micropipette Puller, Sutter Instrument, Novato, CA); the glass pipette provided mechanical rigidity that allowed advancing the tetrode to the deep-lying PG. Prior to recording, tetrode tips were gold-plated (NanoZ 1.4, Multi Channel Systems, Reutlingen, Germany) to obtain 200–300 kOhm impedance at 1kHz. The electrode was positioned above the brain according to stereotaxic brain atlas coordinates (Maler et al., 1991) - 150-300 µm caudal to T26 and 800-1000 µm lateral to midline, and lowered using a micropositioner (Model 2662, David Kopf Instruments, Tujunga, CA) while delivering visual and electrosensory stimuli. Tectal responses to such stimuli (Bastian, 1982) were usually detected twice, around 1200 µm and around 1900 µm ventral to the top of cerebellum (as expected from the curved shape of OT). The electrode usually then transversed nucleus electrosensorius, producing weak multi-unit responses to electrocommunication stimuli around 2300-2500 µm (Heiligenberg et al., 1991). In the initial experiments the location of PG was verified by preparing histological sections and locating the electrode track marks (Wallach et al., 2018). PG units were usually encountered between 2800 µm and 3400 µm ventral to the top of the cerebellum, and were easily identified due to their characteristic rapid spike bursts (Wallach et al., 2018). Differential extracellular voltage was obtained by using one stereotrode/tritrode channel as reference. This enabled near-complete cancellation of the electric organ discharge (EOD) interference.

We report on the responses of 98 PG neurons responsive to electrocommunication signals, within the medial subdivision of PG (PGm); we also found in the same sites 33 cells responding to moving objects, 17 cells responding to passive electrosensory stimuli (ampullary receptors, Bell & Maler, 2005), 5 responding to acoustic stimuli and 5 to visual stimuli. We found no evident topography or spatially organized distribution of the various responses recorded in PG.

The EOD rate of the fish was continuously recorded through a pair of carbon rods places next to the skin. A sine-wave stimulus was delivered via another pair of carbon electrodes via a stimulus isolation unit (A-M systems 2200). The stimulus frequency was set relative to the fish’s EOD frequency to maintain a constant beat pattern. The stimulation intensity was set so that the maximal amplitude at the EOD electrodes would be 10%. In the beat-scan protocol (**Figs. 6 and 7**), beats were delivered in 1 s cycles, each with a randomly set ΔF, in which the stimulus amplitude was sinusoidally modulated to produce a varying envelope signal (see **Fig. 6S1**). Small chirps were delivered by transiently increasing the stimulus frequency by 80 Hz for 14 ms. Long chirps were delivered by transiently increasing the stimulus frequency by 600 Hz and decreasing the stimulus amplitude by 50% for 30 ms. Chirps were delivered in blocks of 30 repetitions, usually at 1 s intervals, on the background of a constant beat stimulus of various ΔFs (small chirps: ΔF=-60 Hz in N=60 cells, ΔF=+5 Hz in N=42 cells, ΔF=+150 Hz in N=7 cells, ΔF=-150 Hz in N=7 cells; long chirps: ΔF=-120 Hz in N=35 cells). In the chirp-scan protocol (**Fig. 8**), a chirp was delivered at the peak of the envelope signal of the beat-scan protocol.

### Quantification and statistical analysis

#### Preprocessing

Data were acquired using Spike2 (Cambridge Electronic Designs, Cambridge, UK) and analyzed in Matlab (MathWorks, Natick, MA). Single units were sorted offline by performing principal component and clustering analyses. Units were considered to be well-separated and to represent individual neurons, only if (a) their spike shapes were homogenous over time and did not overlap with other units or noise; and (b) the unit exhibited a refractory period of >1 ms in autocorrelation histograms (Wallach et al., 2018).

#### Chirp response analysis

Self-chirps were detected off-line by identifying EOD rate excursions exceeding five standard-deviations. Timestamps of exogenous chirp delivery onset were recorded for off-line analysis. For each cell, the following procedure was executed for every chirp type. First, the firing rate was calculated in 7.5 ms bins both inside and outside a response window (100 ms before to 150 ms after chirp onset) for all repetitions. Next, the control samples (the rates in the bins outside the response window) were used to generate a distribution of control peri-stimuli response rates (10^4^ random sets of N samples, where N is the number of chirp repetitions). The PSTH threshold θ_PSTH_ was set at the 99^th^ percentile of the resulting distribution. The cell was considered to be responding to the chirp if the PSTH (average firing rate across all repetitions) exceeded this threshold and spikes were recorded in at least six repetitions. Finally, the first response spike was located in each repetition within a time window where the PSTH exceeded θ_PSTH_. The median of all first spike times, relative to the chirp onset, was defined as the first spike latency of the chirp response.

#### Beat and envelope response analysis

The following procedure was conducted for every cell probed with the beat-scan protocol. The total number of spikes was computed in each cycle of the scan. The resulting spike-counts were sorted according to the cycles’ ΔFs and Lowess smoothing (25% span) was used to produce the beat-frequency tuning of the spike count response, *r*_*beat*_(Δ*F*). To determine the response type, we also computed the symmetrical tuning curve, *r*_*beat*_(|Δ*F*|), by sorting the spike counts according to the absolute-valued ΔF and performing Lowess smoothing (25% span). For some cells, the cycles were interleaved with 1-2 s stimulation gaps; the firing rate in these gaps was used to compute the baseline activity for these cells. For all other cells, the 40^th^ percentile of *r*_*beat*_(|Δ*F*|) was defined as the baseline.

The beat filtering type of each cell was determined as follows. First, we checked if the cell was *baseband permitting*, i.e. if *r*_*beat*_ (|Δ*F*|) surpasses the baseline in at least one of the two cycles closest to ΔF=0. All baseband permitting cells belonged to one of the following 3 types: if 70% of *r*_*beat*_(|Δ*F*|) was above baseline, the cell was defined as APF; if the peak of *r*_*beat*_ (Δ*F*) (the asymmetric tuning curve) was at |ΔF|>50 Hz, the cell was defined as ΔF+ (single side-band filter); otherwise the cell was defined as LPF. For cells that were not baseband permitting, those for which *r*_*beat*_ (|Δ*F*|) exceeded baseline at the maximal |ΔF| were defined as HPF, otherwise they were defined as BPF. The response band for each of the types was defined as the region in which *r*_*beat*_(|Δ*F*|) exceeded baseline. Next, we computed the envelope tuning of each cell, *r*_*env*_(*φ*) where *φ* is the envelope phase, i.e. the time from the beginning of the cycle normalized by the total cycle duration. The envelope tuning was computed using only cycles in which ΔF was within the response band of the cell, and was smoothed using the Lowess method (10% span).

Joint beat-envelope tuning was approximated as the product of two univariate tuning curves: *r*_*beat,evn*_(Δ*F, φ*) = r_*beat*_(Δ*F*) · r_$,-_(*φ*). The response regions (**Fig. 7C**) were defined using the contour where the response was at 50% of the maximum. To estimate the population beat-envelope tuning (**Fig. 7D)**, we averaged all normalized bivariate tuning functions.

### Electric potential and chirp simulations

Simulations for the electric potential of two interacting fish were implemented in the finite-element modelling software COMSOL Multiphysics as two-dimensional electrostatic boundary-value problems. Two fish models, each of which consists of a data-calibrated geometry, skin layer properties, and electric organ, are introduced in a 3 m wide simulation domain where a fine triangular mesh is generated (**Fig. 5S1 panel A**). The current density associated with the EOD is specified as a sum of two Gaussian functions whose parameters are calibrated from data (Babineau et al., 2006) – see **Fig.5S1 Panel B**). The Poisson equation (a partial differential equation whose solution gives the spatial profile of the electric potential) is then solved and yields a dipole-like EOD consistent with observations. Temporally, the EOD waveform is assumed to be purely sinusoidal, and the various phases of the cycle are solved through a parameter sweep of the time variable.

Chirps are implemented as Gaussian frequency excursion, *f*_*chirp*_(*t*), with a maximum of 80 Hz and duration of *Δt* = 15 ms (**Fig.5S1 Panels C and D**). The EOD phase of the chirping fish is thus given by 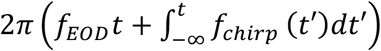 (Benda et al., 2005), where *f*_*EOD*_ is the base EOD frequency. The electric potential of the chirping fish is summed with that of the non-chirping fish, resulting in a beat that is perturbed by the presence of the chirp. Notably, the beat phase exhibits a shift, or reset, when compared to the linear advance expected when no chirps occur, *ΔfΔt*. To evaluate this phase reset, the beat pattern is first extracted from the summed electric potential as the amplitude of its analytic signal (using the Hilbert transform; red curve, **Fig. 5S1 Panel E**). The same approach is then applied to the beat itself in order to obtain its phase,*φ*_*B*_(*t*) (**Fig. 5S1 Panel F**). The phase reset is defined as the beat phase difference between the chirp offset and onset, minus the underlying chirp-independent phase advance: *Δφ*_*C*_= *φ*_*B*_(*Δ t*/2) − *φ*_*B*_(−*Δ t*/2) − Δ*fΔt*, and is evaluated along the length of the fish, from its mid-line to the head (**Fig. 5B**). The tail region is populated by very few electroreceptors (Carr et al., 1982) and therefore does not contribute to perception and was omitted from analysis.

## Supplementary Material

**Table 3S1:**
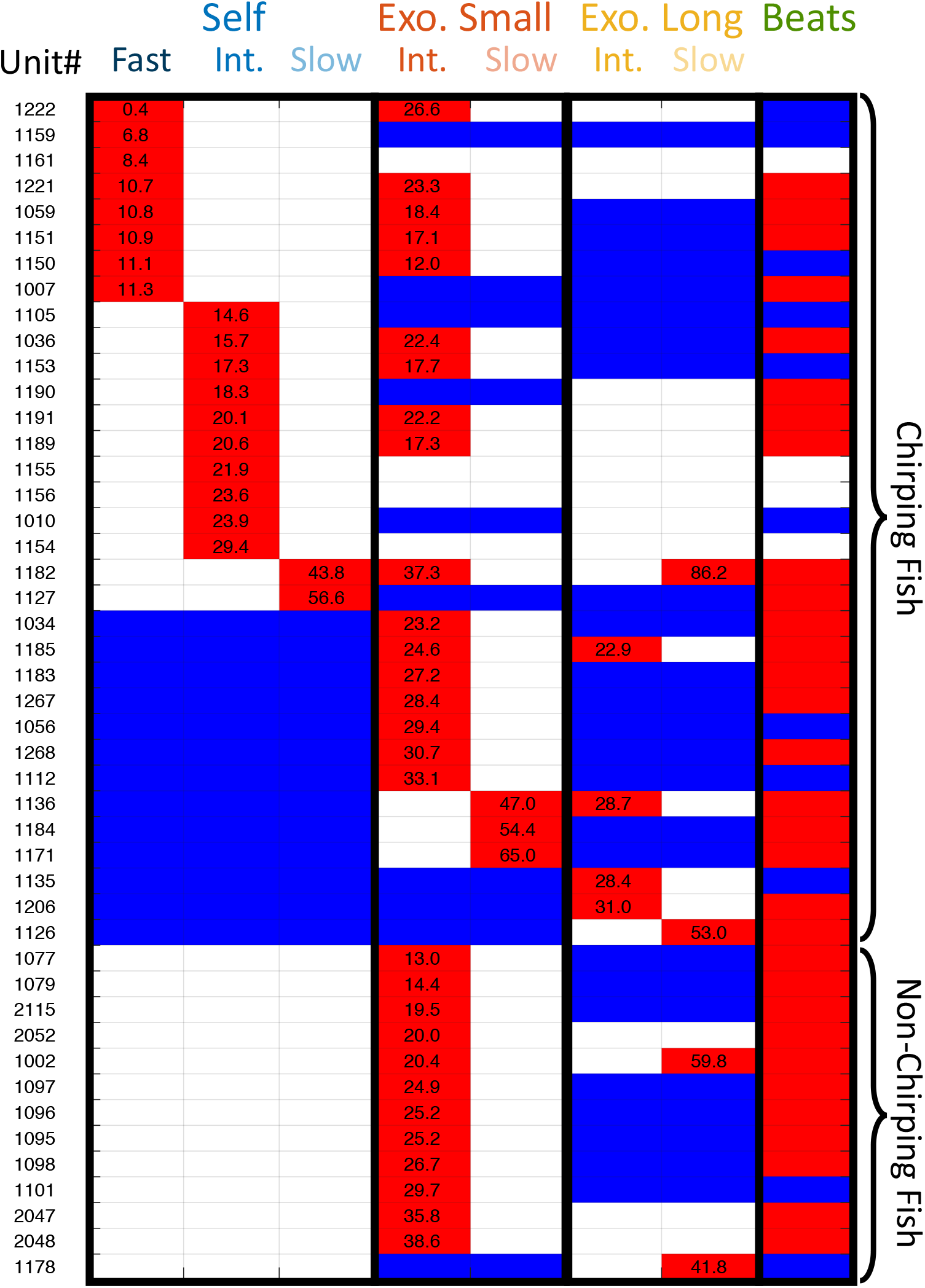
All PG chirp responses. Red: cell responded (number cites median latency); blue: cell did not respond; white: unknown (self: fish did not emit chirps; exogenous: chirp stimuli were not delivered).

**Figure 4S1:**
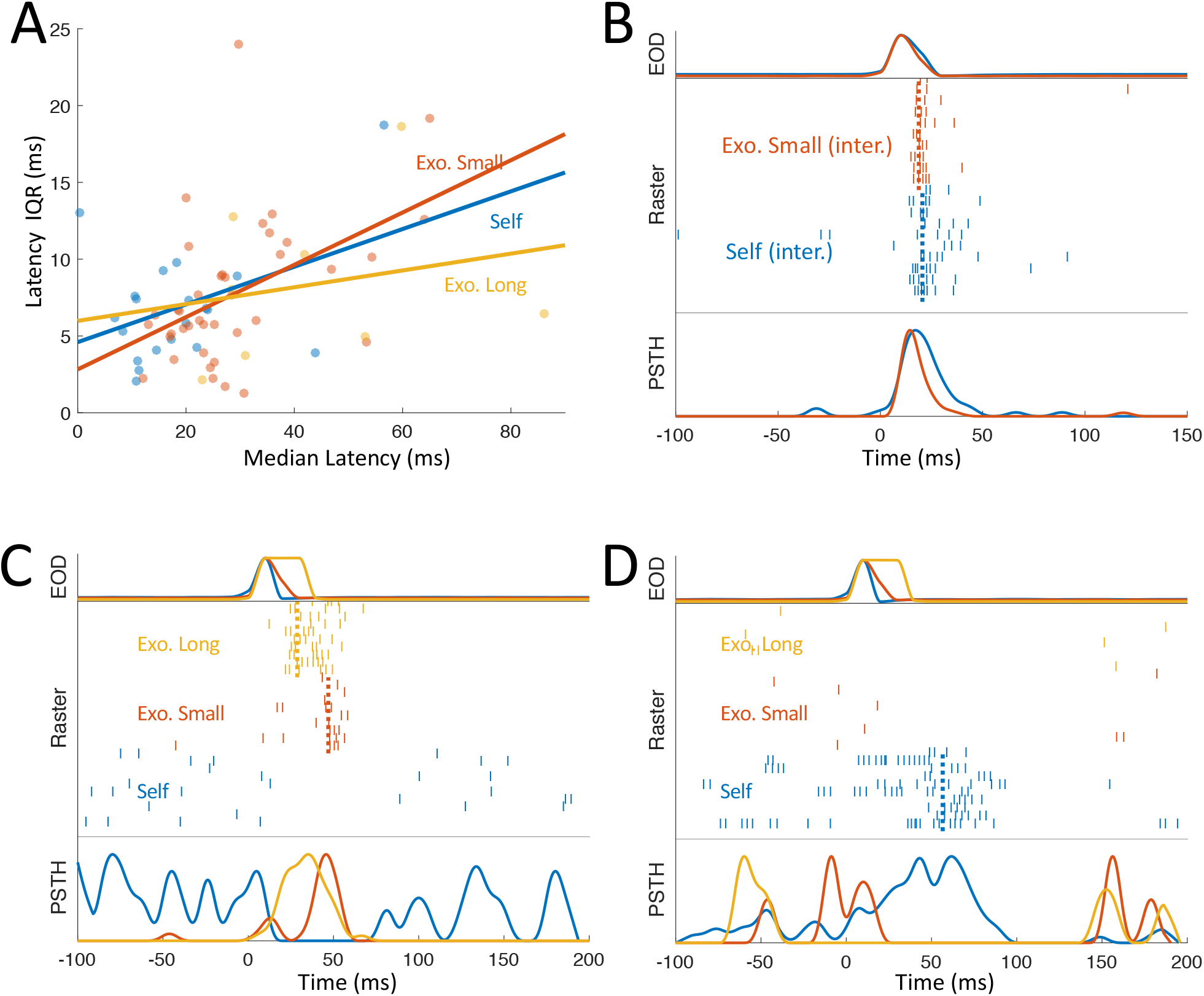
Complex interactions between chirp types in PG. (**A**) For all chirp types, longer median latencies were associated with higher latency variability (lines depict linear regression), suggesting that long latencies reflect longer multi-synaptic pathways. **(B-D)** Top: EOD rate excursion (normalized); Middle: raster plot depicting spike response of one PG cell to individual chirp instances of different kinds (dashed lines-median first spike latency); Bottom: PSTH across chirps. (**B**) A cell responding to self and exogenous chirp with similar latency and pattern, suggesting a common afferent pathway (similar to **Fig. 4D**). (**C**) An example PG cell excited by both small and long exogenous chirps and inhibited by self-chirp. (**D**) An example of a cell excited by self-chirp and inhibited by exogenous chirps.

**Figure 5S1:**
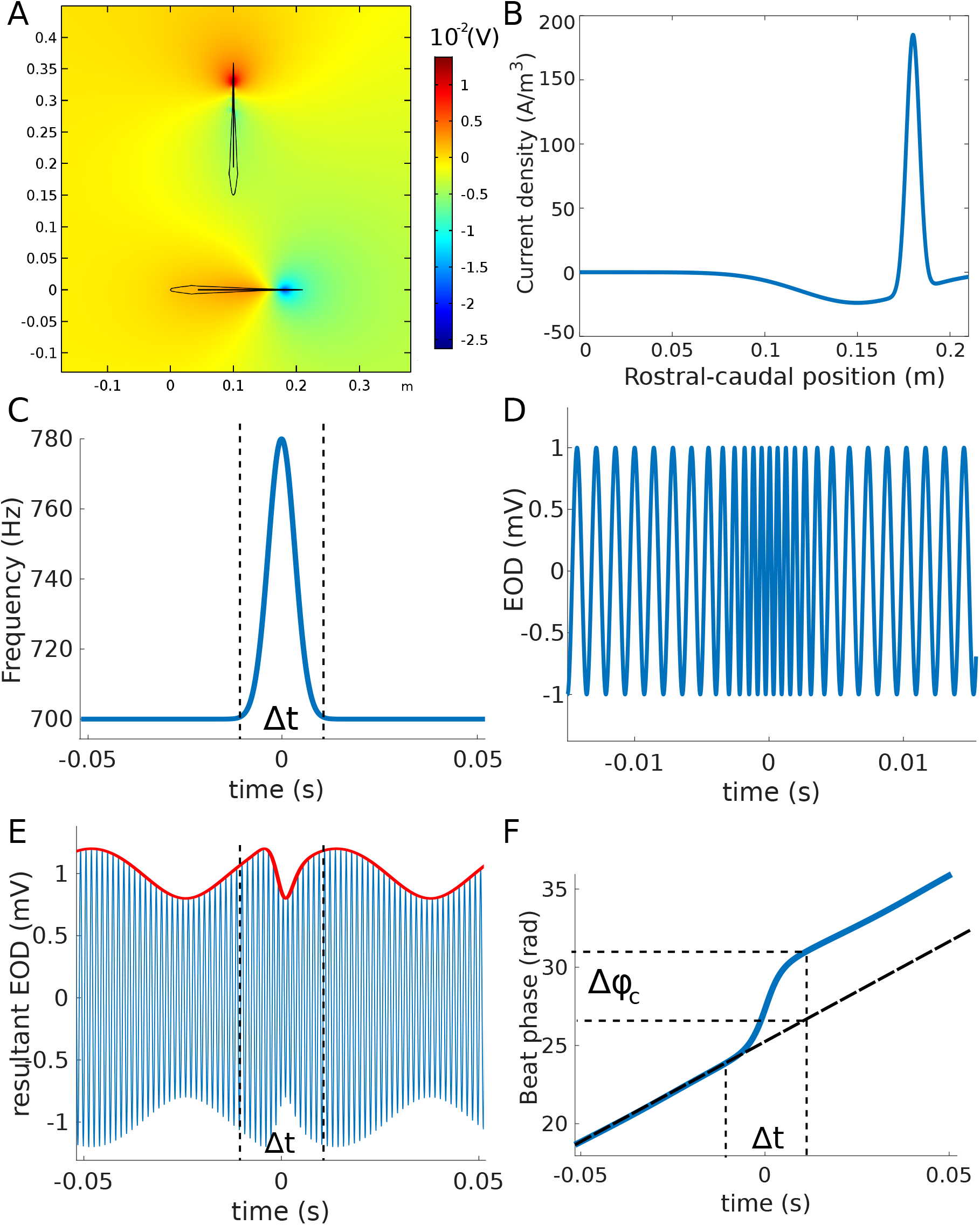
COMSOL Multiphysics simulation. **(A)** Snapshot of the electric potential due to two interacting fish. (**B**) Data-calibrated EOD current source density used in (A). (**C**) Chirps are implemented as a Gaussian frequency excursion. (**D**) Chirp leads to an accelerated sinusoidal EOD trace for the chirping fish. (**E)** When summed with the other fish’s EOD, the chirp affects the the beat amplitude modulation pattern (red curve). (**F**) Beat phase *Δφ*_*C*_without (dashed black) and with (blue) chirp; chirp induces a shift or reset of the beat phase.

**Figure 6S1:**
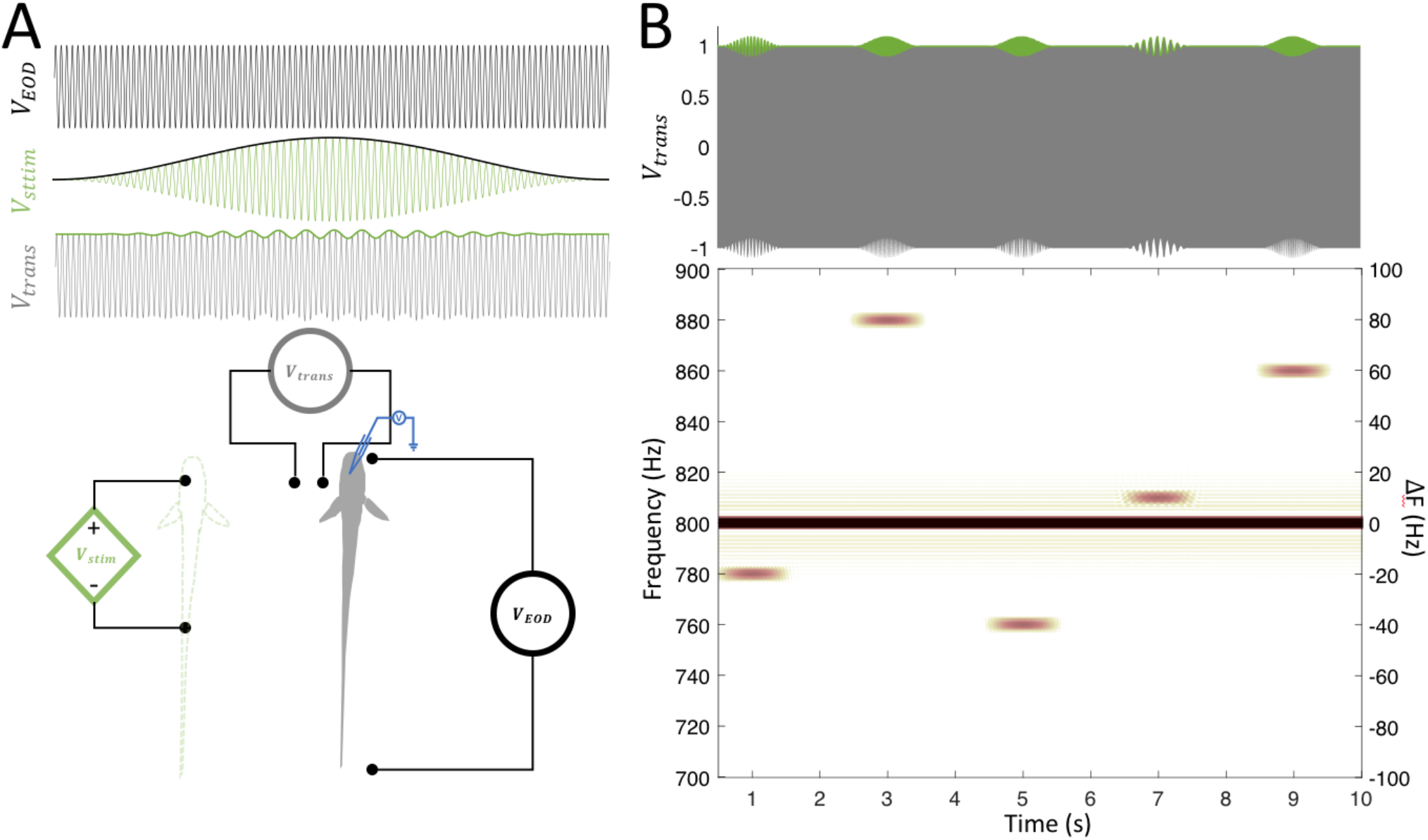
Beat-scan protocol. (**A**) An immobilized fish is placed in the experimental tank. The fishes EOD remains intact and can be monitored (*V*_*EOD*_, black, top). A stimulator is used to deliver sinusoidal wave (*V*_*stim*,_ middle, green) at a set frequency difference from that of the recorded fish, with a sinusoidal envelope (dark green), imitating a conspecific of a certain identity approaching and then retreating; the transdermal potential, approximately measured close to the skin (*V*_*trans*,_ bottom, grey), exhibits a beat pattern with a fixed frequency and varying contrast. (**B**) Top: transdermal potential showing a sequence of several beat cycles, each at a randomly selected beat frequency. Bottom: spectrogram of the transdermal potential.

## References

Allen, K. M., & Marsat, G. (2018, Mar 27). Task-specific sensory coding strategies are matched to detection and discrimination performance. J Exp Biol, 221(Pt 6). https://doi.org/10.1242/jeb.170563

Babineau, D., Longtin, A., & Lewis, J. E. (2006, Sep). Modeling the electric field of weakly electric fish. J Exp Biol, 209(Pt 18), 3636–3651. http://www.ncbi.nlm.nih.gov/entrez/query.fcgi?cmd=Retrieve&db=PubMed&dopt=Citation&list_uids=16943504

Bastian, J. (1982). Vision and electroreception. Integration of sensory information in the optic tectum of the weakly electric fish Apteronotus leptorhynchus. Journal of Comparative Physiology A-Sensory Neural & Behavioral Physiology, 147, 287–297.

Bastian, J., Schneiderjen, S., & Nguyenkim, J. (2001). Arginine vasotocin modulates a sexually dimorphic communication behavior in the weakly electric fish, Apteronotus leptorhynchus. Journal of Experimental Biology, 204, 1909–1923.

Behrend, K., & Donicht, M. (1990). Descending connections from the brainstem to the spinal cord in the electric fish Eigenmannia. Brain Behavior and Evolution, 35, 227–239.

Bell, C., & Maler, L. (2005). Central neuroanatomy of electrosensory systems in fish. In T.H. Bullock & C. Hopkins (Eds.), Electroreception (Vol. 21, pp. 68–111). Springer.

Benda, J., Longtin, A., & Maler, L. (2005, Mar 2). Spike-frequency adaptation separates transient communication signals from background oscillations. J Neurosci, 25(9), 2312–2321. http://www.ncbi.nlm.nih.gov/entrez/query.fcgi?cmd=Retrieve&db=PubMed&dopt=Citation&list_uids=15745957

Benda, J., Longtin, A., & Maler, L. (2006, Oct 19). A synchronization-desynchronization code for natural communication signals. Neuron, 52(2), 347–358. http://www.ncbi.nlm.nih.gov/entrez/query.fcgi?cmd=Retrieve&db=PubMed&dopt=Citation&list_uids=17046696

Berman, N., Dunn, R. J., & Maler, L. (2001). Function of NMDA receptors and persistent sodium channels in a feedback fathway of the electrosensory system. Journal of Neurophysiology, 86, 1612–1621.

Bottai, D., Dunn, R., Ellis, W., & Maler, L. (1997). N-Methyl-D-Aspartate receptor 1 mRNA distribution in the central nervous system of the weakly electric fish Apteronotus leptorhynchus. Journal of Comparative Neurology, 389, 65–80.

Carr, C. E., & Maler, L. (1985). A Golgi study of the cell types of the dorsal torus semicircularis of the electric fish Eigenmannia: functional and morphological diversity in the midbrain. Journal of Comparative Neurology, 235(2), 207–240.

Carr, C. E., Maler, L., Heiligenberg, W., & Sas, E. (1981). Laminar organization of the afferent and efferent systems of the torus semicircularis of gymnotiform fish: Morphological substrates for parallel processing in the electrosensory system. J Comp Neurol, 203, 649–670.

Carr, C. E., Maler, L., & Sas, E. (1982). Peripheral organization and central projections of the electrosensory organs in gymnotiform fish. Journal of Comparative Neurology, 211, 139–153.

Cenquizca, L. A., & Swanson, L. W. (2006, Jul 1). Analysis of direct hippocampal cortical field CA1 axonal projections to diencephalon in the rat. J Comp Neurol, 497(1), 101–114. https://doi.org/10.1002/cne.20985

Chacron, M. J., Longtin, A., & Maler, L. (2011, Oct). Efficient computation via sparse coding in electrosensory neural networks. Current Opinion in Neurobiology, 21(5), 752–760. https://doi.org/10.1016/j.conb.2011.05.016

Chacron, M. J., Maler, L., & Bastian, J. (2005, Apr 3). Electroreceptor neuron dynamics shape information transmission. Nat Neurosci, 8, 673–678. http://www.ncbi.nlm.nih.gov/entrez/query.fcgi?cmd=Retrieve&db=PubMed&dopt=Citation&list_uids=15806098

Clarke, S. E., Longtin, A., & Maler, L. (2015, Dec). Contrast coding in the electrosensory system: parallels with visual computation. Nat Rev Neurosci, 16(12), 733–744. https://doi.org/10.1038/nrn4037

Clarke, S. E., & Maler, L. (2017, May 8). Feedback Synthesizes Neural Codes for Motion. Curr Biol, 27(9), 1356–1361. https://doi.org/10.1016/j.cub.2017.03.068

Cuddy, M., Aubin-Horth, N., & Krahe, R. (2012, Jan). Electrocommunication behaviour and non invasively-measured androgen changes following induced seasonal breeding in the weakly electric fish, Apteronotus leptorhynchus. Hormones and behavior, 61(1), 4–11. https://doi.org/10.1016/j.yhbeh.2011.09.003

Dulka, J. G., & Ebling, S. L. (1999, Apr 24). Testosterone increases the number of substance P-like immunoreactive neurons in a specific sub-division of the lateral hypothalamus of the weakly electric, brown ghost knifefish, Apteronotus leptorhynchus. Brain Res, 826(1), 1–9. https://www.ncbi.nlm.nih.gov/pubmed/10216191

Dulka, J. G., Maler, L., & Ellis, W. (1995). Androgen-induced changes in electrocommunicatory behavior are correlated with changes in substance P-like immunoreactivity in the brain of the electric fish Apteronotus leptorhynchus. Journal of Neuroscience, 15, 1879–1890.

Dunlap, K., & Larkins-Ford, J. (2003). Production of aggressive electrocommunication signals to progressively realistic social stimuli in male Apteronotus leptorhynchus. Ethology, 109, 243–258.

Dunlap, K., Pelczar, P., & Knapp, R. (2002, Sep). Social Interactions and Cortisol Treatment Increase the Production of Aggressive Electrocommunication Signals in Male Electric Fish, Apteronotus leptorhynchus. Hormones and behavior, 42(2), 97. http://www.ncbi.nlm.nih.gov/entrez/query.fcgi?cmd=Retrieve&db=PubMed&dopt=Citation&list_uids=12367563

Dunlap, K. D., & Larkins-Ford, J. (2003, Feb). Diversity in the structure of electrocommunication signals within a genus of electric fish, Apteronotus. J Comp Physiol A Neuroethol Sens Neural Behav Physiol, 189(2), 153–161. http://www.ncbi.nlm.nih.gov/entrez/query.fcgi?cmd=Retrieve&db=PubMed&dopt=Citation&list_uids=12607044

Elliott, S. B., Harvey-Girard, E., Giassi, A. C., & Maler, L. (2017, Jun 13). Hippocampal-like circuitry in the pallium of an electric fish: Possible substrates for recursive pattern separation and completion. J Comp Neurol, 525, 8–46. https://doi.org/10.1002/cne.24060

Elliott, S. B., & Maler, L. (2015, Aug 5). Stimulus induced Up states in the dorsal pallium of a weakly electric fish. J Neurophysiol, 114, 2071–2076. https://doi.org/10.1152/jn.00666.2015

Engler, G., Fogarty, C. M., Banks, J. R., & Zupanc, G. K. H. (2000). Spontaneous modulations of the electric organ discharge in the weakly electric fish, Apteronotus leptorhynchus: a biophysical and behavioral analysis. Journal of Comparative Physiology A, 7/8, 645–660.

Engler, G., & Zupanc, G. K. H. (2001). Differential production of chirping behavior evoked by electrical stimulation of the weakly electric fish, Apteronotus leptorhynchus. Journal of Comparative Physiology A, 187, 274–256.

Fotowat, H., Harrison, R. R., & Krahe, R. (2013, Aug 21). Statistics of the Electrosensory Input in the Freely Swimming Weakly Electric Fish Apteronotus leptorhynchus. J Neurosci, 33(34), 13758–13772. https://doi.org/10.1523/JNEUROSCI.0998-13.2013

Fugere, V., Ortega, H., & Krahe, R. (2011, Apr 23). Electrical signalling of dominance in a wild population of electric fish. Biol Lett, 7(2), 197–200. https://doi.org/10.1098/rsbl.2010.0804

Gao, S. C., Wei, Y. C., Wang, S. R., & Xu, X. H. (2019, Aug). Medial Preoptic Area Modulates Courtship Ultrasonic Vocalization in Adult Male Mice. Neurosci Bull, 35(4), 697–708. https://doi.org/10.1007/s12264-019-00365-w

Giassi, A. C., Correa, S. A., & Hoffmann, A. (2007, Aug 10). Fiber connections of the diencephalic nucleus tuberis anterior in the weakly electric fish, Gymnotus cf. carapo: an in vivo tract-tracing study. J Comp Neurol, 503(5), 655–667. https://doi.org/10.1002/cne.21413

Giassi, A. C., Duarte, T. T., Ellis, W., & Maler, L. (2012). The Organization of the Gymnotiform Fish Pallium in Relation to Learning and Memory: II. Extrinsic connections. Journal of Comparative Neurology, 520, 3338–3368.

Giassi, A. C., Ellis, W., & Maler, L. (2012). The Organization of the Gymnotiform Fish Pallium in Relation to Learning and Memory: III. Intrinsic connections. Journal of Comparative Neurology, 520 3369–3394.

Hagedorn, M., & Heiligenberg, W. (1985). Court and spark: Electric signals in the courtship and mating behavior of gymnotid fish. Animal Behavior, 33, 254–265.

Harvey-Girard, E., Dunn, R. J., & Maler, L. (2007, Dec 20). Regulated expression of N-methyl-D-aspartate receptors and associated proteins in teleost electrosensory system and telencephalon. J Comp Neurol, 505(6), 644–668. https://doi.org/10.1002/cne.21521

Harvey-Girard, E., Giassi, A. C., Ellis, W., & Maler, L. (2012). The Organization of the Gymnotiform Fish Pallium in Relation to Learning and Memory: IV. Expression of Conserved Transcription Factors and Implications for the Evolution of Dorsal Telencephalon. Journal of Comparative Neurology, 520, 3395–3413.

Harvey-Girard, E., Giassi, A. C., Ellis, W., & Maler, L. (2013, Mar 1). Expression of the cannabinoid CB1 receptor in the gymnotiform fish brain and its implications for the organization of the teleost pallium [Research Support, Non-U.S. Gov’t]. J Comp Neurol, 521(4), 949–975. https://doi.org/10.1002/cne.23212

Harvey-Girard, E., Tweedle, J., Ironstone, J., Cuddy, M., Ellis, W., & Maler, L. (2010, Jul 15). Long-term recognition memory of individual conspecifics is associated with telencephalic expression of Egr-1 in the electric fish Apteronotus leptorhynchus. J Comp Neurol, 518(14), 2666–2692. https://doi.org/10.1002/cne.22358

Hashikawa, K., Hashikawa, Y., Tremblay, R., Zhang, J., Feng, J. E., Sabol, A., Piper, W. T., Lee, H., Rudy, B., & Lin, D. (2017, Nov). Esr1(+) cells in the ventromedial hypothalamus control female aggression. Nat Neurosci, 20(11), 1580–1590. https://doi.org/10.1038/nn.4644

Hashikawa, Y., Hashikawa, K., Falkner, A. L., & Lin, D. (2017). Ventromedial Hypothalamus and the Generation of Aggression. Front Syst Neurosci, 11, 94. https://doi.org/10.3389/fnsys.2017.00094

Heiligenberg, W., Keller, C. H., Metzner, W., & Kawasaki, M. (1991). Structure and function of neurons in the complex of the nucleus electrosensorius of the gymnotiform fish Eigenmannia: Detection and processing of electric signals in social communication. Journal of Comparative Physiology A-Sensory Neural & Behavioral Physiology, 169, 151–164.

Heiligenberg, W., Metzner, W., Wong, C. J. H., & Keller, C. H. (1996). Motor control of the jamming avoidance response of Apteronotus leptorhynchus: evolutionary changes of a behavior and its neuronal substrates. Journal of Comparative Physiology A-Sensory Neural & Behavioral Physiology, 179, 653–674.

Heiligenberg, W., & Rose, G. J. (1987, Jul). The optic tectum of the gymnotiform electric fish, Eigenmannia: labeling of physiologically identified cells. Neuroscience, 22(1), 331–340. http://www.ncbi.nlm.nih.gov/entrez/query.fcgi?cmd=Retrieve&db=PubMed&dopt=Citation&list_uids=3627446

Henninger, J., Krahe, R., Kirschbaum, F., Grewe, J., & Benda, J. (2018, May 7). Statistics of natural communication signals observed in the wild identify important yet neglected stimulus regimes in weakly electric fish. J Neurosci. https://doi.org/10.1523/JNEUROSCI.0350-18.2018

Huang, C. G., & Chacron, M. J. (2016, Sep 21). Optimized Parallel Coding of Second-Order Stimulus Features by Heterogeneous Neural Populations. J Neurosci, 36(38), 9859–9872. https://doi.org/10.1523/JNEUROSCI.1433-16.2016

Huang, C. G., Metzen, M. G., & Chacron, M. J. (2018, Oct 5). Feedback optimizes neural coding and perception of natural stimuli. Elife, 7. https://doi.org/10.7554/eLife.38935

Huang, C. G., Metzen, M. G., & Chacron, M. J. (2019, Oct). Descending pathways mediate adaptive optimized coding of natural stimuli in weakly electric fish. Sci Adv, 5(10), eaax2211. https://doi.org/10.1126/sciadv.aax2211

Hupe, G. J., & Lewis, J. E. (2008, May). Electrocommunication signals in free swimming brown ghost knifefish, Apteronotus leptorhynchus. J Exp Biol, 211(Pt 10), 1657–1667. https://doi.org/211/10/1657[pii]10.1242/jeb.013516

Johnston, W. J., Palmer, S. E., & Freedman, D. J. (2020, Feb). Nonlinear mixed selectivity supports reliable neural computation. PLoS Comput Biol, 16(2), e1007544. https://doi.org/10.1371/journal.pcbi.1007544

Jun, J. J., Longtin, A., & Maler, L. (2016). Active sensing associated with spatial learning reveals memory-based attention in an electric fish. Journal of Neurophysiology, 115, 2577– 2592. https://doi.org/10.1152/jn.00979.2015

Kawasaki, M., Maler, L., Rose, G., & Heiligenberg, W. (1988). Anatomical and functional organization of the prepacemaker nucleus in gymnotiform electric fish: the accomodation of two behaviors in one nucleus. Journal of Comparative Neurology, 276, 113–131.

Keller, C. H., Maler, L., & Heiligenberg, W. (1990). Structural and functional organization of a diencephalic sensory-motor interface in the gymnotiform fish, Eigenmannia. Journal of Comparative Neurology, 293, 347–376.

Kirouac, G. J. (2015, Sep). Placing the paraventricular nucleus of the thalamus within the brain circuits that control behavior. Neurosci Biobehav Rev, 56, 315–329. https://doi.org/10.1016/j.neubiorev.2015.08.005

Kishi, T., Tsumori, T., Ono, K., Yokota, S., Ishino, H., & Yasui, Y. (2000, Apr 3). Topographical organization of projections from the subiculum to the hypothalamus in the rat. J Comp Neurol, 419(2), 205–222. https://doi.org/10.1002/(sici)1096-9861(20000403)419:2<205::aid-cne5>3.0.co;2-0

Krahe, R., Bastian, J. A., & Chacron, M. J. (2008, May 28). Temporal processing across multiple topographic maps in the electrosensory system. Journal of Neurophysiology, 100, 852–867. https://doi.org/90300.2008 [pii]10.1152/jn.90300.2008

Krahe, R., & Maler, L. (2014, Feb). Neural maps in the electrosensory system of weakly electric fish [Review]. Curr Opin Neurobiol, 24C, 13–21. https://doi.org/10.1016/j.conb.2013.08.013

Lo, L., Yao, S., Kim, D. W., Cetin, A., Harris, J., Zeng, H., Anderson, D. J., & Weissbourd, B. (2019, Apr 9). Connectional architecture of a mouse hypothalamic circuit node controlling social behavior. Proc Natl Acad Sci U S A, 116(15), 7503–7512. https://doi.org/10.1073/pnas.1817503116

Maler, L. (1979). The posterior lateral line lobe of certain gymnotiform fish. Quantitative light microscopy. Journal of Comparative Neurology, 183, 323–363.

Maler, L., & Ellis, W. G. (1987). Inter-male aggressive signals in weakly electric fish are modulated by monoamines. Behavioural Brain Research, 25, 75–81.

Maler, L., & Monaghan, D. (1991). The distribution of excitatory amino acid binding sites in the brain of an electric fish, Apteronotus leptorhynchus. J Chem Neuroanat, 4, 39–61.

Maler, L., Sas, E., Johnston, S., & Ellis, W. (1991). An atlas of the brain of the weakly electric fish Apteronotus Leptorhynchus. J Chem Neuroanat, 4, 1–38.

Marsat, G., Longtin, A., & Maler, L. (2012, Feb 10). Cellular and circuit properties supporting different sensory coding strategies in electric fish and other systems. Current Opinion in Neurobiology, 22, 1–7. https://doi.org/10.1016/j.conb.2012.01.009

Marsat, G., & Maler, L. (2010, Jul 14). Neural heterogeneity and efficient population codes for communication signals. J Neurophysiol, 104, 2543–2555. https://doi.org/jn.00256.2010 [pii]10.1152/jn.00256.2010

Marsat, G., Proville, R., & Maler, L. (2009). Transient signals trigger synchronous bursts in an identified population of neurons. Journal of Neurophysiology, 102, 714–723.

McGillivray, P., Vonderschen, K., Fortune, E. S., & Chacron, M. J. (2012, Apr 18). Parallel coding of first-and second-order stimulus attributes by midbrain electrosensory neurons. The Journal of neuroscience : the official journal of the Society for Neuroscience, 32(16), 5510–5524. https://doi.org/10.1523/JNEUROSCI.0478-12.2012

Metzen, M. G., & Chacron, M. J. (2019). Envelope Coding and Processing: Implications for Perception and Behavior. In B.A. Carlson, J.A. Sisneros, A. Popper, & R. Fay (Eds.), Electroreception: Fundamental Insights from Comparative Approaches (Vol. 70, pp. 251–278). Springer Nature.

Metzen, M. G., Huang, C. G., & Chacron, M. J. (2018, Jun 25). Descending pathways generate perception of and neural responses to weak sensory input. PLoS Biol, 16(6), e2005239. https://doi.org/10.1371/journal.pbio.2005239

Metzner, W. (1999, May). Neural circuitry for communication and jamming avoidance in gymnotiform electric fish. J Exp Biol, 202 (Pt 10), 1365–1375. http://www.ncbi.nlm.nih.gov/entrez/query.fcgi?cmd=Retrieve&db=PubMed&dopt=Citation&list_uids=10210677

Middleton, J. W., Longtin, A., Benda, J., & Maler, L. (2006, Sep 26). The cellular basis for parallel neural transmission of a high-frequency stimulus and its low-frequency envelope. Proc Natl Acad Sci U S A, 103(39), 14596–14601. http://www.ncbi.nlm.nih.gov/entrez/query.fcgi?cmd=Retrieve&db=PubMed&dopt=Citation&list_uids=16983081

Mueller, T., & Guo, S. (2009, Oct 20). The distribution of GAD67-mRNA in the adult zebrafish (teleost) forebrain reveals a prosomeric pattern and suggests previously unidentified homologies to tetrapods. J Comp Neurol, 516(6), 553–568. https://doi.org/10.1002/cne.22122

Pouso, P., Cabana, A., Goodson, J. L., & Silva, A. (2019). Preoptic Area Activation and Vasotocin Involvement in the Reproductive Behavior of a Weakly Pulse-Type Electric Fish, Brachyhypopomus gauderio. Front Integr Neurosci, 13, 37. https://doi.org/10.3389/fnint.2019.00037

Pouso, P., Radmilovich, M., & Silva, A. (2017, Apr). An immunohistochemical study on the distribution of vasotocin neurons in the brain of two weakly electric fish, Gymnotus omarorum and Brachyhypopomus gauderio. Tissue Cell, 49(2 Pt B), 257–269. https://doi.org/10.1016/j.tice.2017.02.003

Richards, S., & Maler, L. (1996). The distribution of met-enkephalin like immunoreactivity in the brain of Apteronotus leptorhynchus, with emphasis on the electrosensory system. J Chem Neuroanat, 11, 173–190.

Rigotti, M., Barak, O., Warden, M. R., Wang, X. J., Daw, N. D., Miller, E. K., & Fusi, S. (2013, May 19). The importance of mixed selectivity in complex cognitive tasks. Nature, 497(7451), 585–590. https://doi.org/10.1038/nature12160

Rigotti, M., Ben Dayan Rubin, D., Wang, X. J., & Fusi, S. (2010). Internal representation of task rules by recurrent dynamics: the importance of the diversity of neural responses. Front Comput Neurosci, 4, 24. https://doi.org/10.3389/fncom.2010.00024

Santana, U. J., Roque-da-Silva, A. C., Duarte, T. T., & Correa, S. A. (2001, Dec). Interference with the GABAergic system in the dorsolateral telencephalon and modulation of the electric organ discharge frequency in the weakly electric fish Gymnotus carapo. J Comp Physiol A Neuroethol Sens Neural Behav Physiol, 187(11), 925–933. http://www.ncbi.nlm.nih.gov/pubmed/11866190

Saper, C. B. (2000). Hypothalamic connections with the cerebral cortex. Progress in brain research, 126, 39–48. https://doi.org/10.1016/S0079-6123(00)26005-6

Sas, E., & Maler, L. (1986). The optic tectum of gymnotiform teleosts Eigenmannia viriscens and Apteronotus leptorhynchus: a golgi study. Neuroscience, 18, 215–246.

Sas, E., & Maler, L. (1991). Somatostatin-like immunoreactivity in the brain of an electric fish (Apteronotus leptorhynchus) identified with monoclonal antibodies. J Chem Neuroanat, 4(3), 155–186.

Sas, E., Maler, L., & Tinner, B. (1990, Feb 1). Catecholaminergic systems in the brain of a gymnotiform teleost fish: an immunohistochemical study. J Comp Neurol, 292(1), 127–162. https://doi.org/10.1002/cne.902920109

Savard, M., Krahe, R., & Chacron, M. J. (2011, Jan 13). Neural heterogeneities influence envelope and temporal coding at the sensory periphery. Neuroscience, 172, 270–284. https://doi.org/10.1016/j.neuroscience.2010.10.061

Scheich, H., Bullock, T.H., and Hamstra, R.H. (1973). Coding properties of two classes of afferent nerve fibers: high frequency electroreceptors in the electric fish, eigenmania. Journal of Neurophysiology, 36, 39–60.

Stringer, C., Pachitariu, M., Steinmetz, N., Carandini, M., & Harris, K. D. (2019, Jul). High-dimensional geometry of population responses in visual cortex. Nature, 571(7765), 361–365. https://doi.org/10.1038/s41586-019-1346-5

Tallarovic, S. K., & Zakon, H. H. (2002, Sep). Electrocommunication signals in female brown ghost electric knifefish, Apteronotus leptorhynchus. J Comp Physiol A Neuroethol Sens Neural Behav Physiol, 188(8), 649–657. http://www.ncbi.nlm.nih.gov/entrez/query.fcgi?cmd=Retrieve&db=PubMed&dopt=Citation&list_uids=12355241

Telgkamp, P., Combs, N., & Smith, G. T. (2007, Feb 15). Serotonin in a diencephalic nucleus controlling communication in an electric fish: sexual dimorphism and relationship to indicators of dominance. Dev Neurobiol, 67(3), 339–354. http://www.ncbi.nlm.nih.gov/entrez/query.fcgi?cmd=Retrieve&db=PubMed&dopt=Citation&list_uids=17443792

Toscano-Marquez, B., Oboti, L., Harvey-Girard, E., Maler, L., & Krahe, R. (2020, Oct 21). Distribution of the cholinergic nuclei in the brain of the weakly electric fish, Apteronotus leptorhynchus: Implications for sensory processing. J Comp Neurol. https://doi.org/10.1002/cne.25058

Triefenbach, F., & Zakon, H. (2003). Effects of sex, sensitivity and status on cue recognition in the weakly electric fish Apteronotus leptorhynchus. Animal Behavior, 65, 19–28.

Trinh, A. T., Clarke, S. E., Harvey-Girard, E., & Maler, L. (2019, Jul/Aug). Cellular and Network Mechanisms May Generate Sparse Coding of Sequential Object Encounters in Hippocampal-Like Circuits. eNeuro, 6(4). https://doi.org/10.1523/ENEURO.0108-19.2019

Trinh, A. T., Harvey-Girard, E., Teixeira, F., & Maler, L. (2016, Aug 3). Cryptic laminar and columnar organization in the dorsolateral pallium of a weakly electric fish. J Comp Neurol, 524, 408–428. https://doi.org/10.1002/cne.23874

Vertes, R. P., Linley, S. B., & Hoover, W. B. (2015, Jul). Limbic circuitry of the midline thalamus. Neurosci Biobehav Rev, 54, 89–107. https://doi.org/10.1016/j.neubiorev.2015.01.014

Vonderschen, K., & Chacron, M. J. (2011, Dec). Sparse and dense coding of natural stimuli by distinct midbrain neuron subpopulations in weakly electric fish [Research Support, Non-U.S. Gov’t]. Journal of Neurophysiology, 106(6), 3102–3118. https://doi.org/10.1152/jn.00588.2011

Wallach, A., Harvey-Girard, E., Jun, J. J., Longtin, A., & Maler, L. (2018, Nov 22). A time-stamp mechanism may provide temporal information necessary for egocentric to allocentric spatial transformations. Elife, 7, e36769. https://doi.org/10.7554/eLife.36769

Walz, H., Grewe, J., & Benda, J. (2014, Aug 15). Static frequency tuning accounts for changes in neural synchrony evoked by transient communication signals. J Neurophysiol, 112(4), 752–765. https://doi.org/10.1152/jn.00576.2013

Walz, H., Hupe, G. J., Benda, J., & Lewis, J. E. (2012, Sep 5). The neuroethology of electrocommunication: How signal background influences sensory encoding and behaviour in Apteronotus leptorhynchus. J Physiol Paris. https://doi.org/10.1016/j.jphysparis.2012.07.001

Wei, Y. C., Wang, S. R., Jiao, Z. L., Zhang, W., Lin, J. K., Li, X. Y., Li, S. S., Zhang, X., & Xu, X. H. (2018, Jan 18). Medial preoptic area in mice is capable of mediating sexually dimorphic behaviors regardless of gender. Nat Commun, 9(1), 279. https://doi.org/10.1038/s41467-017-02648-0

Weld, M. M., & Maler, L. (1992). Substance P-like immunoreactivity in the brain of the gymnotiform fish Apteronotus leptorhynchus: presence of sex differences. J Chem Neuroanat, 5, 107–129.

Wong, C. J. (2000, Jan). Electrical stimulation of the preoptic area in Eigenmannia: evoked interruptions in the electric organ discharge. J Comp Physiol [A], 186(1), 81–93. http://www.ncbi.nlm.nih.gov/entrez/query.fcgi?cmd=Retrieve&db=PubMed&dopt=Citation&list_uids=10659045

Wong, C. J. H. (1997a). Afferent and efferent connections of the diencephalic prepacemaker nucleus in the weakly electric fish, Eigenmannia viriscens: Interactions between the electromotor system and the neuroendocrine axis. Journal of Comparative Neurology, 383, 18–41.

Wong, C. J. H. (1997b). Connections of the basal forebrain of the weakly electric fish, Eigenmannia viriscens. Journal of Comparative Neurology, 389, 49–64.

Yamamoto, T., Maler, L., & Nagy, J. I. (1992). Organization of galanin-like immunoreactive neuronal systems in weakly electric fish (Apteronotus leptorhynchus). J Chem Neuroanat, 5, 19–38.

Yang, C. F., Chiang, M. C., Gray, D. C., Prabhakaran, M., Alvarado, M., Juntti, S. A., Unger, E. K., Wells, J. A., & Shah, N. M. (2013, May 9). Sexually dimorphic neurons in the ventromedial hypothalamus govern mating in both sexes and aggression in males. Cell, 153(4), 896–909. https://doi.org/10.1016/j.cell.2013.04.017

Yu, N., Hupe, G., Garfinkle, C., Lewis, J. E., & Longtin, A. (2012, Jul). Coding conspecific identity and motion in the electric sense. PLoS Comput Biol, 8(7), e1002564. https://doi.org/10.1371/journal.pcbi.1002564

Yu, N., Hupe, G., Longtin, A., & Lewis, J. E. (2019). Electrosensory Contrast Signals for Interacting Weakly Electric Fish. Front Integr Neurosci, 13, 36. https://doi.org/10.3389/fnint.2019.00036

Zakon, H., Oestreich, J., Tallarovic, S., & Triefenbach, F. (2002, Sep-Dec). EOD modulations of brown ghost electric fish: JARs, chirps, rises, and dips. J Physiol Paris, 96(5-6), 451–458. http://www.ncbi.nlm.nih.gov/entrez/query.fcgi?cmd=Retrieve&db=PubMed&dopt=Citation&list_uids=14692493

Zupanc, G. K. (2002, Sep-Dec). From oscillators to modulators: behavioral and neural control of modulations of the electric organ discharge in the gymnotiform fish, Apteronotus leptorhynchus. J Physiol Paris, 96(5-6), 459–472. http://www.ncbi.nlm.nih.gov/entrez/query.fcgi?cmd=Retrieve&db=PubMed&dopt=Citation&list_uids=14692494

Zupanc, G. K., Sirbulescu, R. F., Nichols, A., & Ilies, I. (2006, Feb). Electric interactions through chirping behavior in the weakly electric fish, Apteronotus leptorhynchus. J Comp Physiol A Neuroethol Sens Neural Behav Physiol, 192(2), 159–173. http://www.ncbi.nlm.nih.gov/entrez/query.fcgi?cmd=Retrieve&db=PubMed&dopt=Citation&list_uids=16247622

Zupanc, G. K. H., & Maler, L. (1993). Evoked chirping in the weakly electric fish Apteronotus leptorhynchus: a quantitative biophysical analysis. Canadian Journal of Zoology, 71, 2301–2310.

